# Acarbose ameliorates Western diet-induced metabolic and cognitive impairments in the 3xTg mouse model of Alzheimer’s disease

**DOI:** 10.1101/2024.06.27.600472

**Authors:** Michelle M. Sonsalla, Reji Babygirija, Madeline Johnson, Samuel Cai, Mari Cole, Chung-Yang Yeh, Isaac Grunow, Yang Liu, Diana Vertein, Mariah F. Calubag, Michaela E. Trautman, Cara L. Green, Michael J. Rigby, Luigi Puglielli, Dudley W. Lamming

## Abstract

Age is the greatest risk factor for Alzheimer’s disease (AD) as well as for other disorders that increase the risk of AD such as diabetes and obesity. There is growing interest in determining if interventions that promote metabolic health can prevent or delay AD. Acarbose is an anti-diabetic drug that not only improves glucose homeostasis, but also extends the lifespan of wild-type mice. Here, we test the hypothesis that acarbose will not only preserve metabolic health, but also slow or prevent AD pathology and cognitive deficits in 3xTg mice, a model of AD, fed either a Control diet or a high-fat, high-sucrose Western diet (WD). We find that acarbose decreases the body weight and adiposity of WD-fed 3xTg mice, increasing energy expenditure while also stimulating food consumption, and improves glycemic control. Both male and female WD-fed 3xTg mice have worsened cognitive deficits than Control-fed mice, and these deficits are ameliorated by acarbose treatment. Molecular and histological analysis of tau and amyloid pathology identified sex-specific effects of acarbose which are uncoupled from the dramatic improvements in cognition, suggesting that the benefits of acarbose on AD are largely driven by improved metabolic health. In conclusion, our results suggest that acarbose may be a promising intervention to prevent, delay, or even treat AD, especially in individuals consuming a Western diet.

## Introduction

As the aging population grows, so does the number of individuals afflicted with age-related diseases, including Alzheimer’s disease (AD). It is estimated that 6.7 million individuals in the United States alone are living with AD, a number that is projected to double by the year 2050 (1, 2). Along with the greatly increasing prevalence of AD are exorbitantly high medical costs. In total, AD and related dementias cost the United States approximately 345 billion dollars in 2023, which is projected to increase to one trillion dollars by the year 2050 (1, 2). Both the high economic cost and prevalence of AD highlight the importance of identifying effective treatment options.

Diabetes and obesity are also significant risk factors for AD (1), and glucose tolerance is known to be impaired in AD patients (reviewed in (3)). As such, factors that contribute to metabolic health, including diet, may play a large role in the development of AD. For years, Western diet (WD), defined as high caloric intake with high levels of carbohydrates, fats (particularly saturated fats), and cholesterol (4–6), has been implicated in the obesity epidemic and the development and progression of T2DM. Consuming a WD just one day a week leads to sustained insulin resistance and fatty liver disease in mice (7). A link has also been established between WD and AD (5, 6, 8–11). Long term consumption of a WD exacerbates many aspects of AD pathology, including neuroinflammation, β-amyloid pathology, and tau pathology in several mouse models of AD (5, 8, 9, 11).

Acarbose is an α-glucosidase inhibitor that has been prescribed for treatment of T2DM since the late 1900’s (12, 13). After a meal, α-glucosidase in the small intestines cleaves polysaccharides, which allows the smaller monosaccharides to be taken up by the gut, resulting in a post-prandial spike in blood glucose levels. When acarbose is used, α-glucosidase activity is blocked, leading to slowed and reduced overall glucose absorption and lowered post-prandial hyperglycemia (12, 13). The benefits of acarbose on metabolic health have been shown in both mice and humans, and include reductions in body weight and improved glucose homeostasis (14–18). Acarbose has also been shown to extend lifespan in genetically heterogenous mice, suggesting that it may act to slow the aging process, especially in males (16, 17).

As acarbose improves metabolic health and slows aging, it might also be able to combat AD through a variety of mechanisms. Acarbose reduces the activity of the mechanistic target of rapamycin complex 1 (mTORC1) in both liver and kidneys (19), and genetic or pharmaceutical inhibition of mTORC1 has been found to extend lifespan in diverse species (reviewed in (20, 21)). mTORC1 signaling is also strongly linked to AD, with human AD patients exhibiting increased mTORC1 activation, and inhibition of mTORC1 with rapamycin has been shown to improve cognition and delay AD pathology in multiple mouse models (22–26). Conversely, acarbose boosts the activity of a second mTOR complex, mTOR complex 2 (mTORC2) in male mice (27). mTORC2 is necessary for normal neuronal morphology and synaptic plasticity, and its activity may be reduced in AD, suggesting a role for decreased mTORC2 in AD pathology (28). mTORC1 and mTORC2 have both been shown to have effects on autophagy, which is dysfunctional in AD patients (29–31). Finally, acarbose has been shown to reduce age-related increases in inflammation in liver and kidney via inhibition of the MEK3/p38/MK2 pathway (32), and to reduce hypothalamic inflammation specifically in males (33).

The triple-transgenic (3xTg) mouse expresses familial human isoforms of APP (APP_Swe_), Tau (tau_P301L_), and Presenilin (PS1_M146V_), which leads it to develop both Aβ and tau pathology as well as cognitive deficits (26, 34–36). The 3xTg mouse exhibits tau phosphorylation by 6 months of age, followed by Aβ in both the hippocampus and cortex by approximately 12 months of age. Cognitive decline is apparent by 6 months of age. Previous work in 3xTg mice utilizing a high-fat diet has shown that consumption of a high-fat diet leads to worsened metabolic health, aggravated amyloid and tau pathology, and impaired cognitive function (37–39).

We hypothesized that a high-fat high-sucrose Western diet (WD) would exacerbate the pathology and symptoms of AD, which acarbose would be effective at ameliorating. Here, we preconditioned 3xTg mice by feeding either Con or WD from 2-6 months of age, inducing obesity and insulin resistance in WD-fed mice. We then initiated acarbose treatment in the diets of both 3xTg and non-transgenic (NTg) controls at 6 months of age, coinciding with the beginnings of cognitive decline in the model, and roughly equivalent to an AD patient first beginning to experience clinical symptoms. We assessed the effects of both WD and acarbose on metabolic health, AD pathology, cognition, and survival.

We find that WD negatively impacts both metabolic health and cognition in 3xTg mice, with sex-dependent effects on AD pathology. Acarbose treatment had minimal effects on metabolic health, cognition, and AD pathology in the context of a low-fat control diet, but when administered to WD-fed 3xTg mice acarbose ameliorated almost all WD-induced impairments of metabolic health and cognition. The effects of a WD and acarbose on mTOR signaling, autophagy, and AD pathology was strongly sex dependent. In females, WD decreased mTORC2 signaling and increased autophagy. In males, mTORC2 signaling was increased by WD and autophagy was decreased. Acarbose reversed WD-induced mTORC2 changes in females and autophagy changes in males. With respect to AD pathology, WD-fed females exhibited a decrease in total tau and insoluble Aβ40. In males, WD increased both total and phosphorylated tau levels as well as insoluble Aβ40, while soluble Aβ40 levels as well as neuroinflammation were decreased by acarbose. WD impairs cognition in both male and female 3xTg mice, and acarbose completely reverses these effects. Finally, we find that WD dramatically increases the mortality of male 3xTg mice, but acarbose treatment is not sufficient to improve survival. In conclusion, our findings suggest that acarbose may have the potential to improve not only metabolic health, but also to preserve cognition in AD patients who are suffering from metabolic syndrome or consuming a WD.

## Results

### Acarbose improves body weight, fat mass, and metabolic health in Western diet-fed mice

We randomly assigned male and female 3xTg mice and non-transgenic (B6129SF2/J; NTg) mice to one of four diet groups: 1) a low-fat Control diet (Con); 2) a Control diet containing 1,000 ppm acarbose (Con+ACA); 3) a high-fat, high-sucrose Western diet (WD); or 4) a WD containing 1,000 ppm acarbose (WD+ACA). WD and WD+ACA mice were preconditioned on WD for 4 months prior to the start of the study (from 2 to 6 months of age); likewise, Con and Con+ACA mice were fed Con in parallel; all mice were then switched to their final diet groups, containing acarbose or not, at 6 months of age. An acarbose concentration of 1,000 ppm was chosen due to the ability of this dose to extend lifespan and improve the healthspan of aging mice (16, 40). The detailed composition of each diet is provided in **Table S1**.

The experimental design is summarized in **Fig. 1A**; in addition to the indicated procedures, we examined mice longitudinally, tracking their body weight monthly and determining body composition at the beginning of the experiment (6 months of age) and at the end of the experiment after 6 months on diet (12 months of age). As expected, both female and male 3xTg mice preconditioned on WD from 2 to 6 months of age had significantly greater body weight, fat mass, and adiposity than their Con-fed counterparts (**Figs. 1B-E, G-J**). Acarbose treatment immediately began reducing the body weight of WD-fed 3xTg mice of both sexes, significantly reducing fat mass and slightly reducing lean mass after 6 months of treatment; the overall effect was one of significantly reduced adiposity. Acarbose also blunted the accretion of body and fat mass and overall adiposity in 3xTg females, but not in males, consuming the low-fat Con diet (**Figs. 1C, E**). These improvements in body weight and composition were not the result of reduced food consumption; instead, acarbose stimulated food consumption in both male and female 3xTg mice, regardless of whether the mice were fed Con or WD (**Figs. 1F, K**). Overall, acarbose administration in the context of WD dramatically reduced body weight and improved body composition despite increased food consumption.

**Figure 1:**
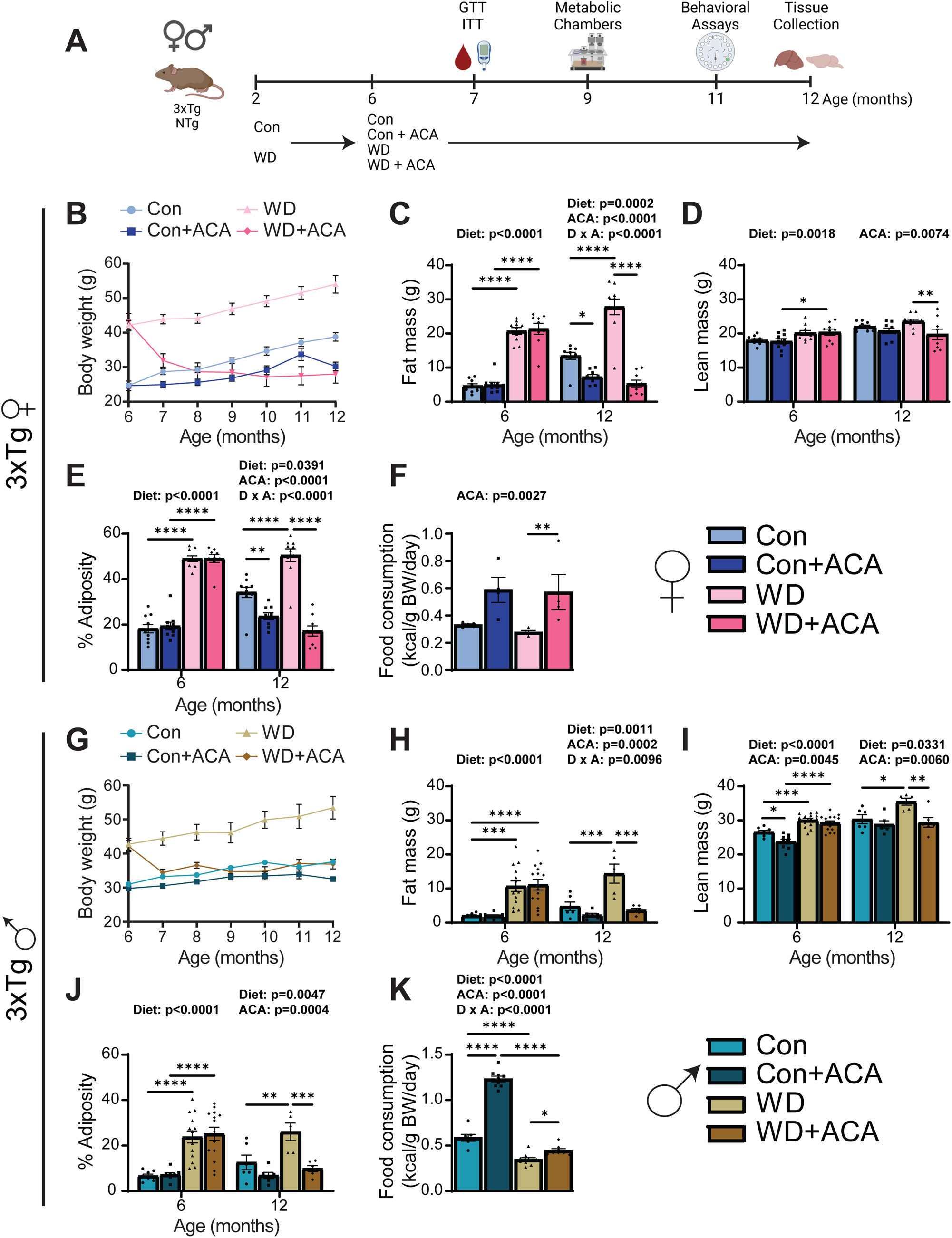
WD-induced changes in the body mass and adiposity of 3xTg mice are ameliorated by acarbose. (A) Experimental design: male and female 3xTg and non-transgenic (NTg) mice were placed on the indicated diets at 2 months of age, then at 6 months of age, half of each group was swapped onto the same diets containing 1,000ppm acarbose; mice periodically underwent metabolic and cognitive phenotyping until 12 months of age. Schematic created with Biorender.com. (B-E) In female 3xTg mice, body weight (B) was measured over the course of the experiment while mice consumed the indicated diets; fat (C) and lean mass (D) were determined at the beginning and end of the experiment, and adiposity (E) was calculated; n = 9-10 mice/group. (F) Food consumption of female 3xTg mice was measured in home cages at 10 months of age and normalized to body weight; n = 4-5 cages/group. (G-J) In male 3xTg mice, body weight (G) was measured over the course of the experiment while mice consumed the indicated diets; fat (H) and lean mass (I) were determined at the beginning and end of the experiment, and adiposity (J) was calculated; n = 5-14 mice/group. (K) Food consumption of male 3xTg mice was measured in home cages at 10 months of age and normalized to body weight; n = 7-10 cages/group. (C-F, H-K) Statistics for the overall effects of diet, acarbose treatment (ACA), and the interaction represent the p-value from a 2-way ANOVA conducted separately for each time point (when applicable), with Sidak post-test, *p < 0.05, **p < 0.01, ***p < 0.001, and ****p < 0.0001. Data represented as mean ± SEM.

NTg females had increased body mass, fat and lean mass, and adiposity when fed a WD, and this was completely ablated by administration of acarbose (**Figs. S1A-D**). Acarbose treatment in Con-fed females did not induce any changes in body mass or composition. Despite the decreased weight of WD-fed NTg females, treatment with acarbose significantly increased food consumption (**Fig. S1E**). Male NTg mice also increased in body mass, fat mass, and adiposity when fed WD, which was reversed by acarbose treatment (**Figs. S1F-I**). Acarbose also decreased the fat mass and adiposity in Con-fed males (**Figs. S1G,I**). Acarbose treatment increased food consumption in Con-fed, but not WD-fed, males (**Fig. S1J**). In summary, acarbose is effective at ameliorating WD-induced changes to body mass and composition in both sexes of 3xTg and NTg mice, with acarbose having minimal effects on Con-fed mice.

As acarbose-treated mice exhibited decreased body mass and reduced fat mass and adiposity despite increased food consumption, we examined energy balance parameters using metabolic chambers. Both female and male 3xTg mice exhibited decreased energy expenditure on WD, which was reversed by acarbose (**Figs. 2A, D**). WD-fed NTg mice exhibited a similar response to acarbose (**Figs. S2A, D**). These differences in energy expenditure were not associated with changes in activity in 3xTg females, but acarbose did increase activity in WD-fed 3xTg males, which reached statistical significance during the light cycle. (**Figs. 2B, E**). In NTg mice, there was a significant decrease in activity during the dark cycle due to WD in females but no alterations in activity in male NTg mice by either diet or treatment (**Figs. S2B, E**). Finally, we assessed substrate utilization by measuring the respiratory exchange ratio (RER), which is calculated from the ratio of O_2_ consumed and CO_2_ released; when the RER approaches 1.0, carbohydrates are primarily being utilized for energy production, and when the RER approaches 0.7, lipids are primarily being utilized (41, 42). As expected, WD-fed mice of both sexes had significantly decreased RER, signifying that WD-fed mice use significantly more lipids for their energy production than carbohydrates; there was no effect of acarbose (**Figs. 2C, F**). NTg males exhibit a similar response to WD feeding, but females have no significant changes to RER in any group (**Figs. S2C, F**). Thus, acarbose may improve body composition in part through increased energy expenditure, which may be due to increased activity in male mice.

**Figure 2:**
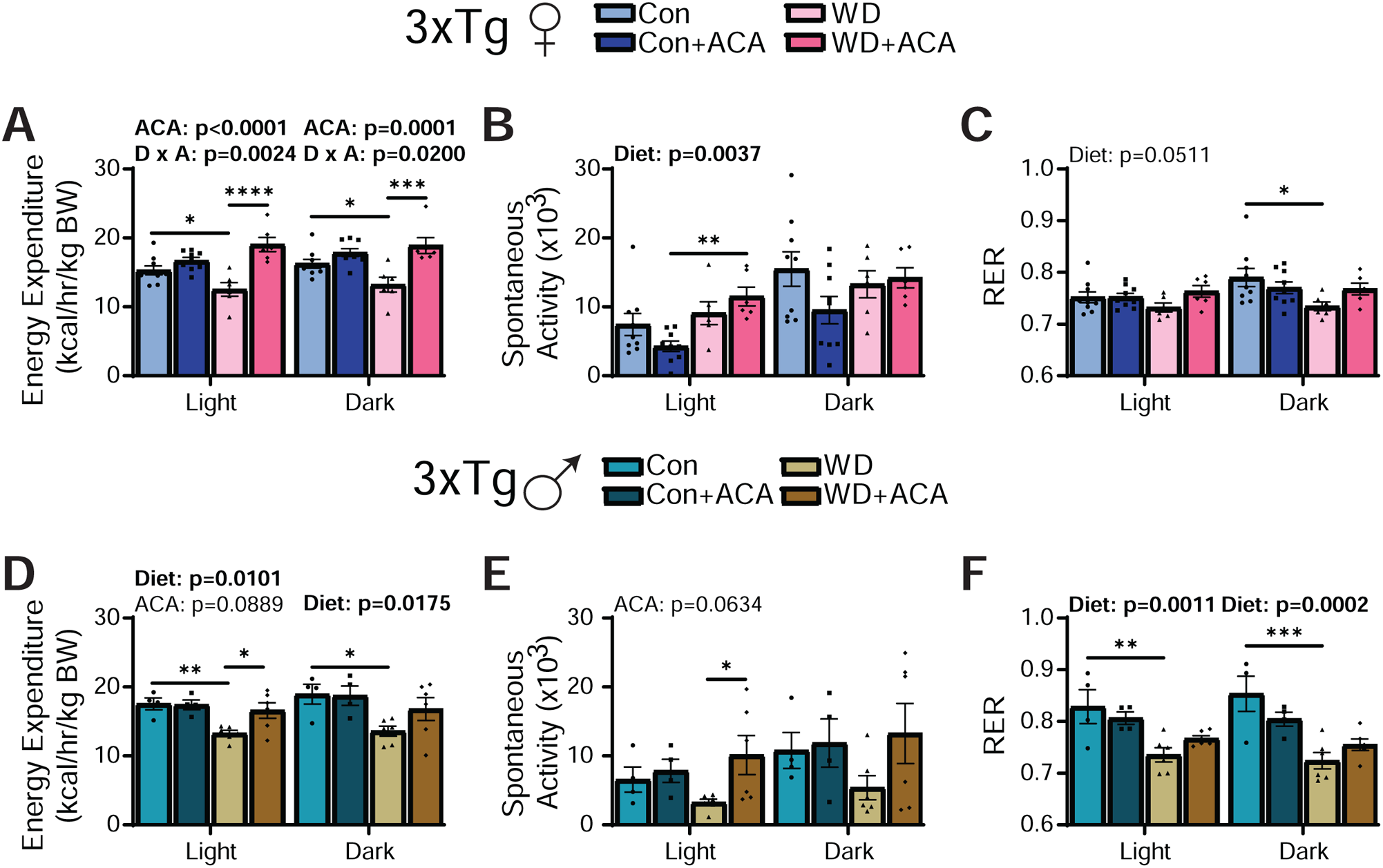
The effects of WD on energy balance in 3xTg mice are opposed by acarbose. (A-F) Metabolic chambers were used to measure energy expenditure, spontaneous activity, and the respiratory exchange ratio (RER) in female (A-C) and male (D-F) 3xTg mice over a 24 hour period. (A, D) Energy expenditure normalized to body weight in females (A) and males (D). (B, E) Spontaneous activity of females (B) and males (E). (C, F) RER of females (C) and males (F). (A-F) n = 4-9 mice/group, statistics for the overall effects of diet, acarbose treatment (ACA), and the interaction represent the p-value from a 2-way ANOVA conducted for the light and dark periods separately, with Sidak post-test, *p < 0.05, **p < 0.01, ***p < 0.001, and ****p < 0.0001. Data represented as mean ± SEM.

Acarbose acts to improve glycemic control by blunting the postprandial blood glucose spike, improving insulin sensitivity in both humans and mice (12). We assessed the effect of both WD and acarbose treatment on glucose tolerance and insulin sensitivity at 7 months of age, after the mice had been on Con or WD diet for 5 months and acarbose treatment for 1 month. In female 3xTg mice, WD feeding for 5 months mildly but significantly exacerbated glucose intolerance, and this effect was reversed by acarbose (**Fig. 3A**). While WD did not significantly impair glucose tolerance in 3xTg males, acarbose was still effective at improving glucose tolerance in WD-fed 3xTg males (**Fig. 3C**). No significant effect on insulin sensitivity was observed with either WD or acarbose in 3xTg mice of either sex (**Figs. 3B, D**). We observed similar effects of WD and acarbose on glucose and insulin tolerance on NTg mice, except that Con-fed NTg males had improved insulin tolerance when treated with acarbose (**Figs. S2G-J**).

**Figure 3:**
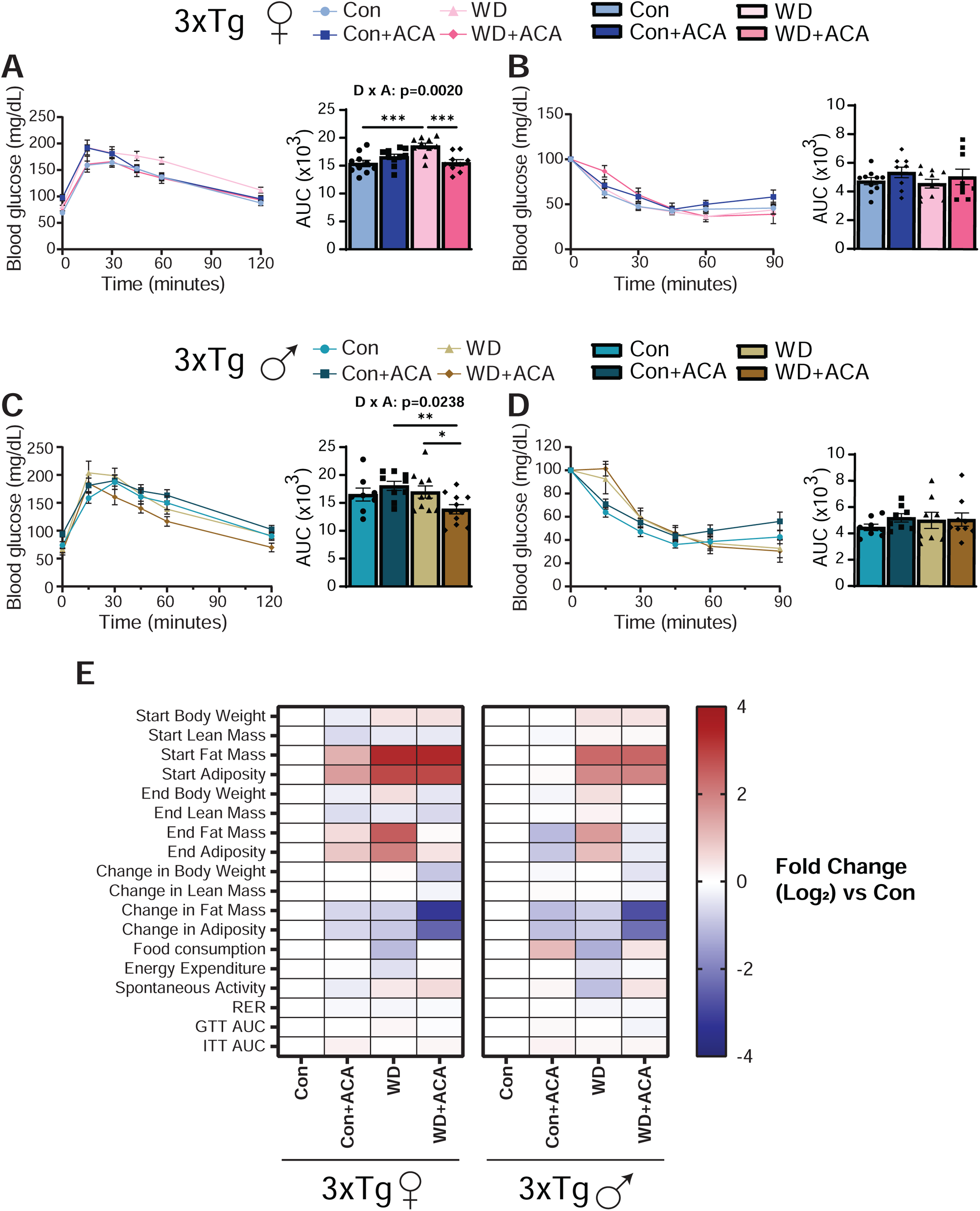
Acarbose improves glucose tolerance in WD-fed 3xTg mice of both sexes. (A-B) Glucose (A) and insulin (B) tolerance tests were performed in female mice at 7 months of age, n = 9-10 mice/group. (C-D) Glucose (C) and insulin (D) tolerance tests were performed in male mice at 7 months of age, n = 8-11 mice/group. (E) Heat map representation of all the metabolic parameters measured in female and male 3xTg mice; color represents the log_2_ fold-change vs. Con-fed 3xTg mice of the same sex. (A-D) Statistics for the overall effects of diet, acarbose treatment (ACA), and the interaction represent the p-value from a 2-way ANOVA, with Sidak post-test, *p < 0.05, **p < 0.01, and ***p < 0.001. Data represented as mean ± SEM.

In summary, WD had expected negative effects on aspects of metabolic health, including weight, adiposity, and glycemic control in both sexes of 3xTg mice and NTg mice, and acarbose ameliorated many of these effects (**Fig. 3E**). As a whole, acarbose did not provide substantial benefits when administered in conjunction with a Con diet aside from a modest improvement in body mass and adiposity.

### Western diet and acarbose induced sex and strain-dependent changes in mTOR signaling and autophagy

mTOR is a master regulator of nutrient signaling, and hyperactivation of mTORC1 has been implicated in the pathology of AD and other age-related diseases (26, 43). Since mTORC1 is activated by nutrient overload, we hypothesized that WD would lead to an increase in mTORC1 activity, which could drive AD pathology. In contrast, acarbose has been shown to inhibit age-related increases in mTORC1 activity (19), and boost mTORC2 activity in male liver (27), which we hypothesized would reduce or delay AD pathology.

We therefore performed Western blotting on protein lysate from whole brains to examine the effect of WD and acarbose on mTORC1 and mTORC2 activity and autophagy (**Figs. 4A-H**). We measured phosphorylation of pS240/244 S6 and pS473 AKT to assess the activity of mTORC1 and mTORC2, respectively. There was an overall effect of WD towards increasing mTORC1 activity (as assessed by phosphorylation of S240/244 S6) in 3xTg females (p=0.0772); there was no significant effect of WD on mTORC1 activity in 3xTg males, and no effect of acarbose on mTORC1 activity in either sex (**Figs. 4A-B, E-F**). In contrast, there was no effect of WD or acarbose on mTORC1 activity in NTg females, however, there was an overall effect of WD, acarbose, and a significant interaction on mTORC1 activity in NTg males, with WD reducing mTORC1 activity, and acarbose significantly reducing mTORC1 activity in Control-fed mice (**Figs. S3A-B, E-F**).

**Figure 4:**
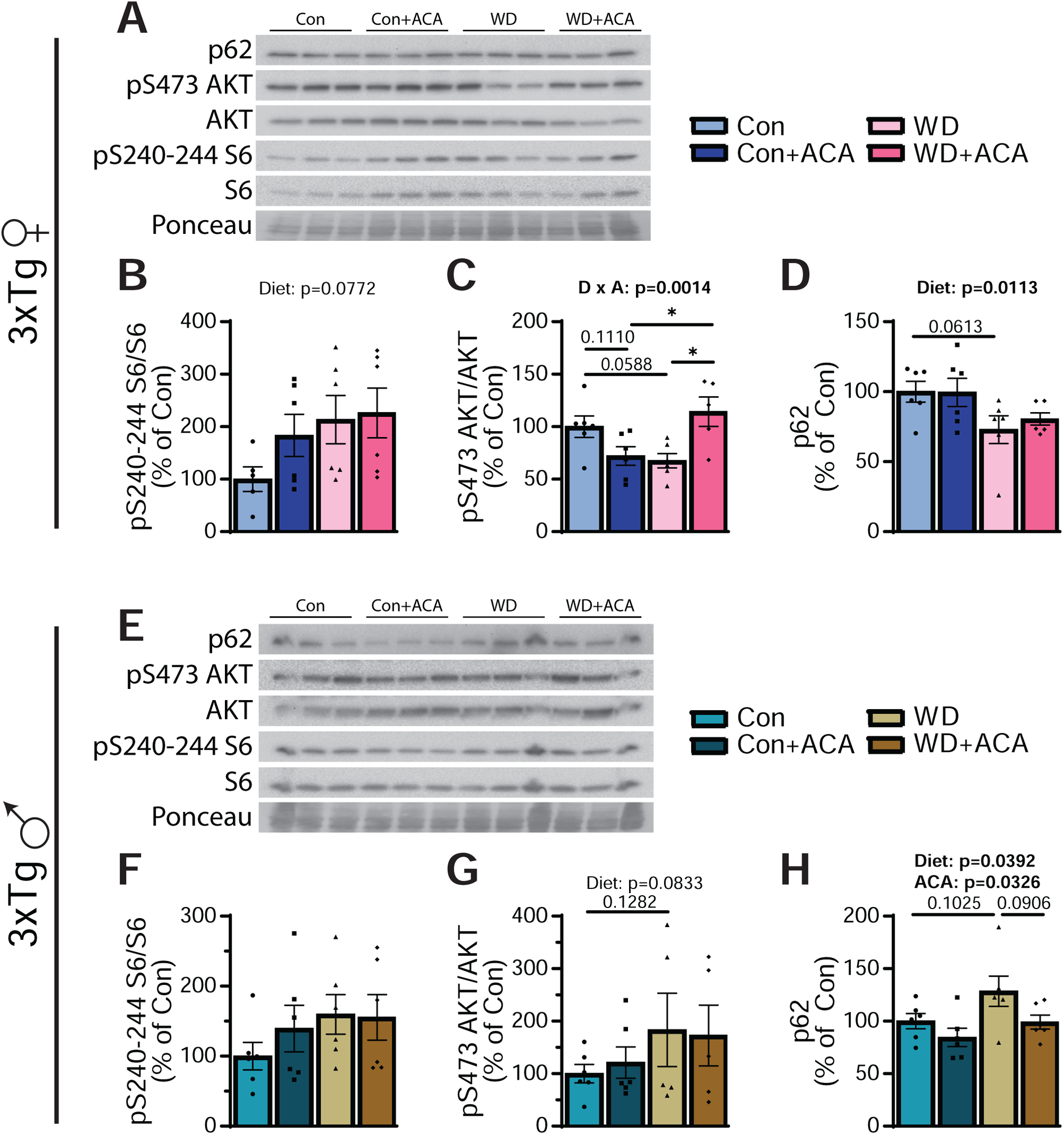
Sex-specific effects of WD and acarbose on mTOR signaling and autophagy. (A-H) mTOR signaling and autophagy were analyzed in the brains of 12-13 month old 3xTg mice via Western blotting of whole brain lysate. (A, E) Representative Western blots of female (A) and male (E) mice. (B, F) Phosphorylation of S240/244 S6 relative to total S6 in female (B) and male (F) mice. (C, G) Phosphorylation of S473 AKT relative to total AKT in female (C) and male (G) mice. (D, H) Quantification of p62 relative to total protein (Ponceau) in female (D) and male (H) mice. (B-D, F-H) n = 5-6 mice/group; statistics for the overall effects of diet, acarbose treatment (ACA), and the interaction represent the p-value from a 2-way ANOVA, with Sidak post-test, *p < 0.05 and **p < 0.01. Data represented as mean ± SEM.

We observed a significant interaction between diet and acarbose on mTORC2 activity (as assessed via S473 AKT phosphorylation), with acarbose significantly increasing mTORC2 activity in WD-fed 3xTg females (**Figs. 4A, C**). In 3xTg males, there was an overall increase in mTORC2 activity on WD (p=0.0833), but the males were not responsive to acarbose (**Figs. 4E, G**). WD increased mTORC2 activity in both female and male NTg mice, but there was no effect of acarbose in either sex (**Figs. S3A, C, E, G**).

Reduced autophagy in AD, which may be partly mediated by abnormal mTOR activity, is a key contributor to the accumulation of plaques and tangles as it disrupts the ability of the brain to clear protein aggregates (44). To assess the effect of WD and acarbose on autophagy, we examined levels of p62 (sequestosome-1; SQSTM1). WD significantly decreased p62 levels in 3xTg females – indicating an increase in autophagy — but there was no effect of acarbose (**Fig. 4A, D**). In 3xTg males WD increased p62 levels – indicating a decrease in autophagy – which was rescued by acarbose (**Figs. 4E, H**). Neither WD nor acarbose significantly affected p62 levels in NTg mice (**Figs. S3A, D, E, H**).

### Western diet and acarbose alter astrocytic, neuronal, and microglial cell populations in a sex-specific manner

In both humans and mice, AD pathology correlates with alterations in the populations of various brain cell types. We assessed GFAP (glial fibrillary acidic protein), NeuN (neuronal nuclei), and Iba1 (ionized calcium binding adaptor 1) as measures of astrocytic, neuronal, and microglial cell populations, respectively, both in the whole brain via Western blotting (**Figs. 5A-H**) and localized in the cortex and hippocampus (via immunofluorescent staining) (**Figs. 6A-H**).

**Figure 5:**
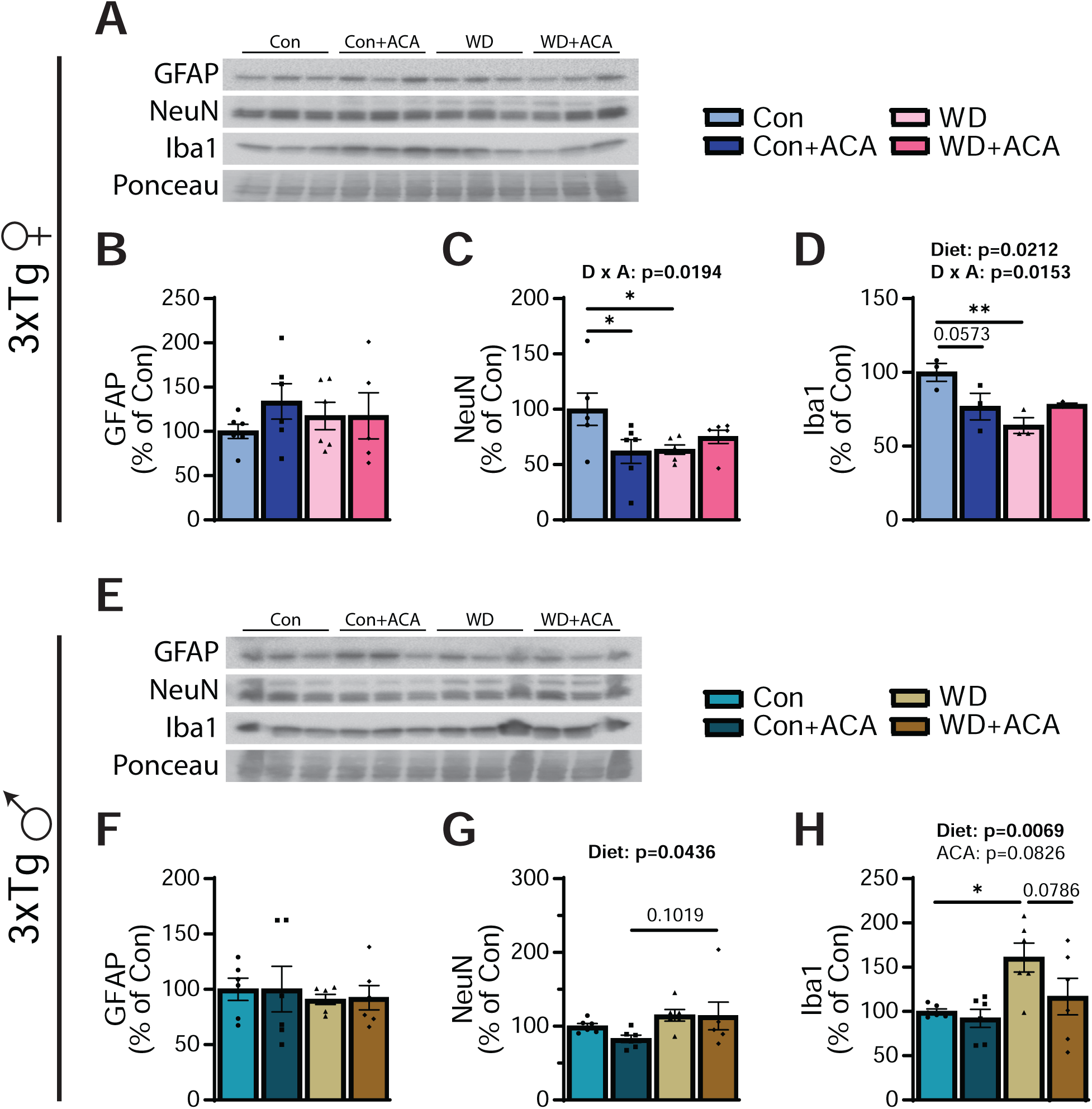
Sex-specific effects of WD and acarbose on NeuN and Iba1. (A-H) Levels of GFAP, NeuN, and Iba1 were analyzed in the brains of 12-13 month old 3xTg mice via Western blotting of whole brain lysate. (A, E) Representative Western blots of female (A) and male (E) mice. (B, F) Quantification of the astrocyte marker GFAP relative to total protein (Ponceau) in female (B) and male (F) mice. (C, G) Quantification of the neuronal marker NeuN relative to total protein (Ponceau) in female (C) and male (G) mice. (D, H) Quantification of the microglial marker Iba1 relative to total protein (Ponceau) in female (D) and male (H) mice. (B-D, F-H) n = 5-6 mice/group; statistics for the overall effects of diet, acarbose treatment (ACA), and the interaction represent the p-value from a 2-way ANOVA, with Sidak post-test, *p < 0.05 and **p < 0.01. Data represented as mean ± SEM.

**Figure 6:**
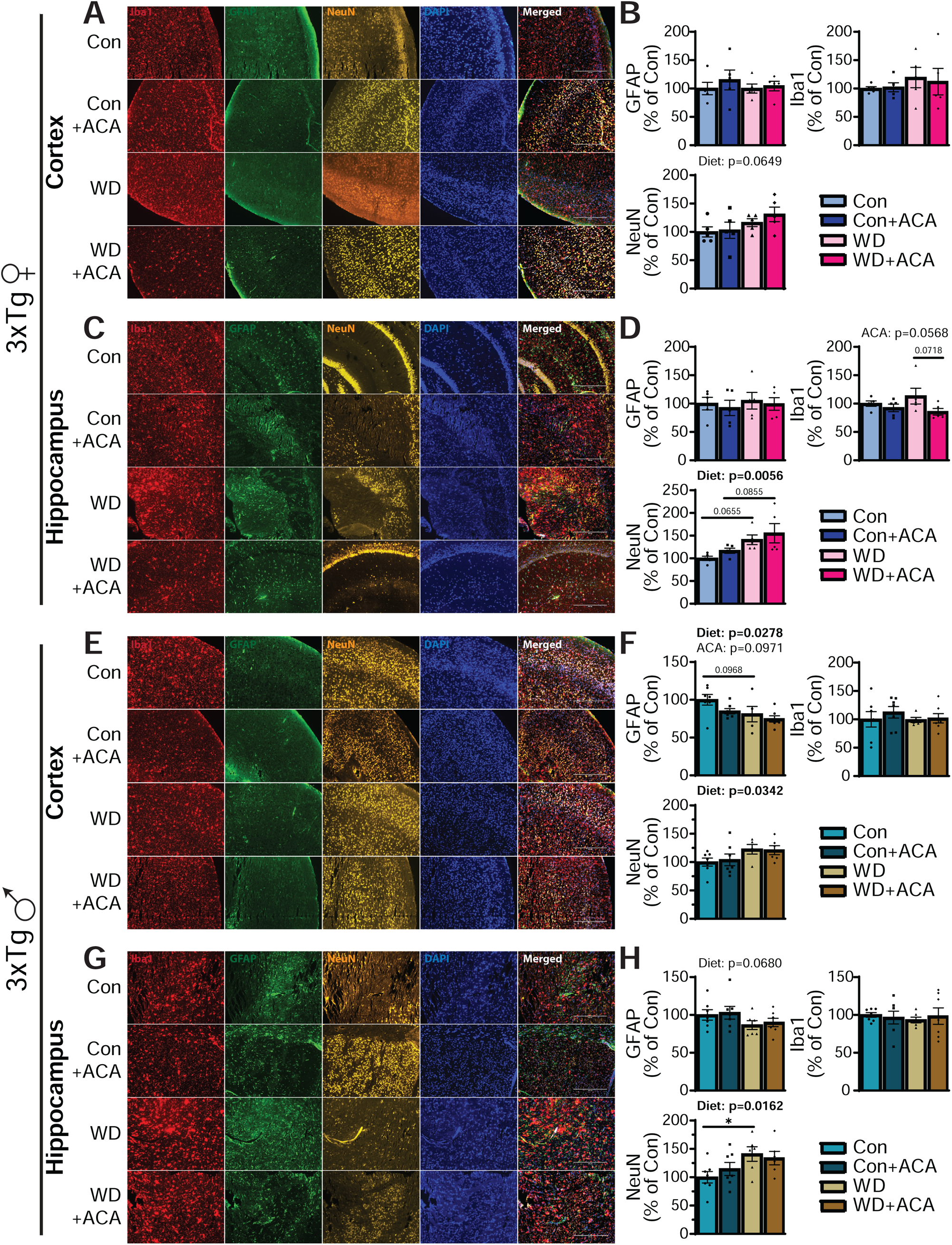
WD and acarbose impact astrocyte, neuron, and microglia populations in the cortex and hippocampus of 3xTg mice. (A, C, E, G) 5 μm paraffin-embedded brain slices were immunostained for GFAP, NeuN, and Iba1 to measure astrocyte, neuron, and microglial populations respectively in the cortex (A, E) and hippocampus (C, G) of 3xTg mice.10x magnification shown; scale bar in image is 400 μm. Results in the cortex and hippocampus are quantified in both female (B, D) and male (F, H) mice. (B, D, F, H) n = 5-6 mice/group; statistics for the overall effects of diet, acarbose treatment (ACA), and the interaction represent the p-value from a 2-way ANOVA, with Sidak post-test, *p < 0.05. Data represented as mean ± SEM.

GFAP is a cytoskeleton intermediate filament protein found in astrocytes that is commonly localized near Aβ plaques and is increased in AD patients and in 3xTg mice (34). We found that GFAP levels were not altered by WD or acarbose in the whole brain, cortex, or hippocampus of 3xTg females (**Figs. 5A-B, and 6A-D**). GFAP levels were decreased in both the cortex and hippocampus in WD-fed 3xTg males (**Figs. 5E-F, and 6E-H**). NeuN is a commonly used marker for assessing neuronal degeneration; we found a sex-specific effect of WD on NeuN levels in the whole brain, with decreased NeuN in WD-fed 3xTg females and increased NeuN in WD-fed 3xTg males; acarbose decreased the level of NeuN in Con-fed 3xTg females (**Figs. 5C, G**). In the cortex and hippocampus, WD increased NeuN levels in both sexes (**Figs 6A-H**). Finally, Iba1 was measured to determine microglial levels. In the whole brain, WD decreased levels of Iba1 in 3xTg females but increased them in 3xTg males; this male-specific increase was rescued by acarbose (p=0.0785) (**Figs. 5D, H**). Hippocampal Iba1 expression was decreased (p = 0.0718) by acarbose treatment in WD-fed 3xTg females; there was no effect in males, although microglia in the hippocampus of WD-fed, non-treated males exhibited a morphologically apparent shift towards more activated microglia (**Figs. 6A-H**).

### Sex-specific response of AD pathology to WD and acarbose in 3xTg mice

We assessed the progression of the two primary hallmarks of AD pathology: the accumulation of phosphorylated tau and Aβ in the brain. We performed Western blotting and an ELISA to assess the progression of AD in the whole brain of 3xTg mice (**Fig. 7**). In female 3xTg mice, we observed that levels of total tau were decreased by both WD and by acarbose (p=0.0532) (**Fig. 7A-B**). However, there was an overall increase in the level phosphorylated tau T231 by acarbose (p=0.0830) and an overall increase of phosphorylated tau T231 (normalized to total tau) with both WD and acarbose (**Figs. 7C-D**). In 3xTg males, WD increased both overall tau levels (p=0.1257) and T231 phosphorylated tau, resulting no overall change in the ratio of phosphorylated to total tau (**Figs. 7H-K**).

**Figure 7:**
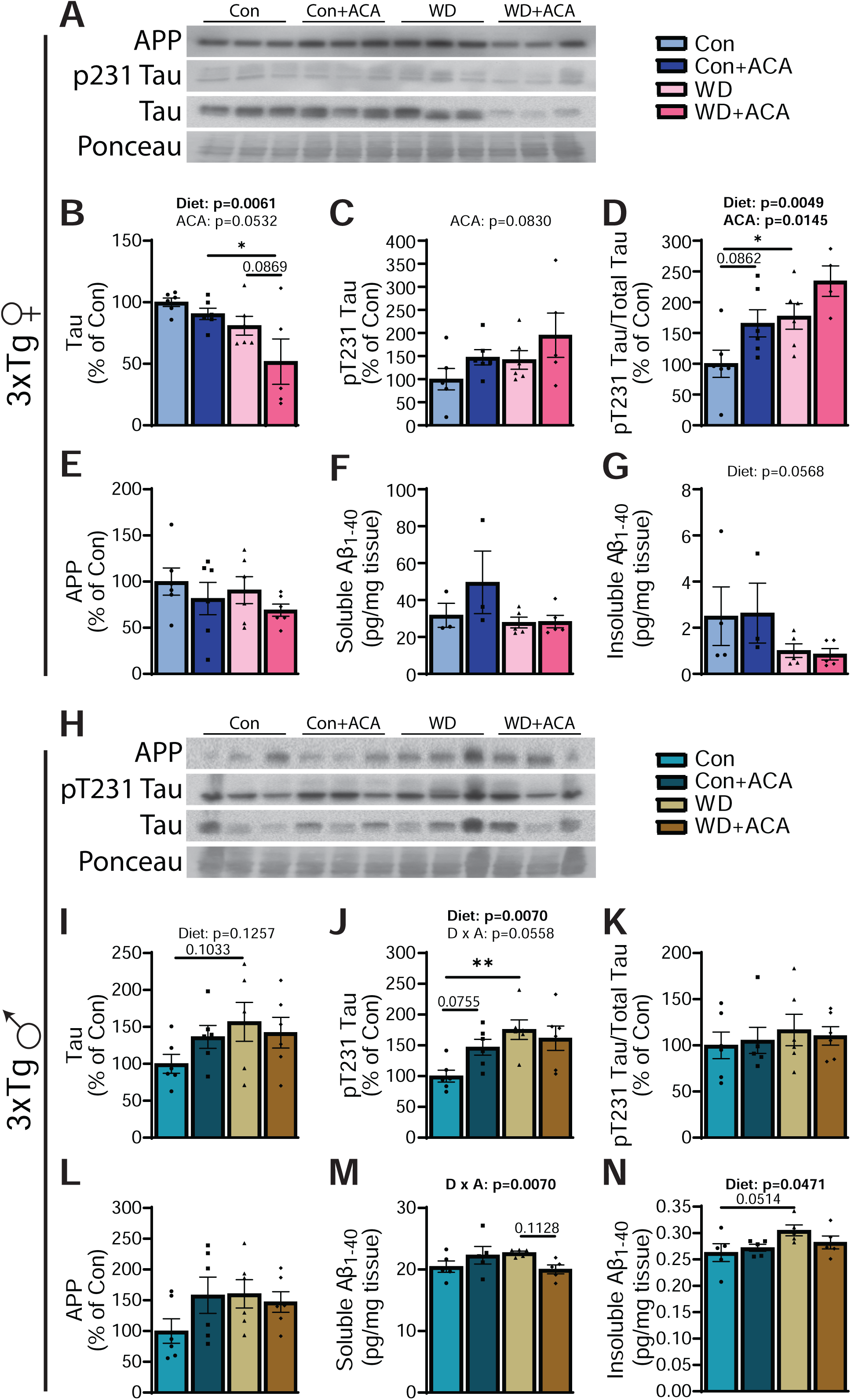
Sex-specific effects of WD and acarbose on AD pathology in 3xTg mice. (A-N) Tau and amyloid pathology was assessed via Western blot and ELISA in female (A-G) and male (H-N) 3xTg mice. (A-G) Representative Western blots from 3xTg female mice are shown in (A) and quantified in (B-E). T231 phosphorylated tau (B) and total tau (C) levels were measured relative to total protein (Ponceau). In addition, the ratio of phosphorylated T231 tau to total tau (D) was calculated. The level of amyloid precursor protein (APP) was measured relative to ponceau staining (E); n = 5-6 mice/group. (F-G) Soluble (F) and insoluble (G) Aβ40 levels were measured by ELISA; n = 4-5 mice/group. (H-L) Representative Western blots from 3xTg male mice are shown in (H) and quantified in (I-L). T231 phosphorylated tau (I) and total tau (J) levels were measured relative to total protein (Ponceau). In addition, the ratio of phosphorylated T231 tau to total tau (K) was calculated. The levels of APP was measured relative to total protein (Ponceau) (L); n = 5-6 mice/group. (M-N) Soluble (M) and insoluble (N) Aβ40 levels were measured by ELISA; n = 4-5 mice/group. (B-G, I-N) Statistics for the overall effects of diet, acarbose treatment (ACA), and the interaction represent the p-value from a 2-way ANOVA, with Sidak post-test, *p < 0.05 and **p < 0.01. Data represented as mean ± SEM

Amyloid precursor protein (APP) and soluble Aβ40 levels were not altered by either WD or acarbose in females 3xTg mice (**Figs. 7E-F**); however, levels of insoluble Aβ40 were lower in WD-fed mice (p=0.0568) (**Fig. 7G**). We also utilized immunohistochemistry to examine Aβ plaque load in the hippocampus and cortex of 3xTg mice but observed minimal Aβ aggregation in any of the diets (**Fig. S4A**). Only 1-2 mice per diet group showed any measurable plaques; of these the mice with the most numerous plaques (via DAB staining as indicated with the arrow in the top leftmost figure) in each diet are shown.

There was no effect of either WD or acarbose on APP levels in 3xTg males (**Fig. 7L**). However, there was an interaction between WD and acarbose on soluble Aβ40 levels (p=0.0070), which we interpret as acarbose reducing soluble Aβ40 only in WD-fed 3xTg males (**Fig. 7M**). WD increased insoluble Aβ40 levels (p=0.0514) but the effect of acarbose was not statistically significant (**Fig. 7N**). Similar to female mice, only 1-2 mice per diet exhibited any plaque accumulation with the most abundant plaque loads shown in **Fig. S4A**.

### Acarbose ameliorates Western diet-induced deficits in spatial learning

To assess the effect of WD and acarbose treatment on cognitive function, we performed behavioral assays at 11-12 months of age. To test spatial learning, we examined performance using Barnes maze (**Figs. 8A-F**). Female 3xTg mice consuming a WD had strongly impaired spatial learning and memory in Barnes maze, as indicated by increased latency to goal during training days 2-4 and during the short and long-term memory tests (**Fig. 8A**) as well as significantly worsened success rate during the same periods (**Fig. 8B**). In addition, WD-fed 3xTg females also exhibited increased latency to any hole during the Barnes maze, indicating worsened anxiety (**Fig. 8C**). Acarbose treatment rescued almost all the WD-induced cognitive deficits in 3xTg females (**Figs. 8A-C**). Male 3xTg mice exhibited similar, though less pronounced, WD-induced deficits in spatial learning, memory, and anxiety from WD; acarbose did not significantly rescue any of these deficits, but there was a trend towards improvement with acarbose with respect to latency to goal and latency to any hole (**Figs. 8D-F**). There were much more limited changes in cognition due to WD or acarbose in NTg mice of either sex (**Figs. S5A-F**), however male WD-fed mice exhibited improved latency to goal and success rate with acarbose treatment (**Figs. S5D-E**).

**Figure 8:**
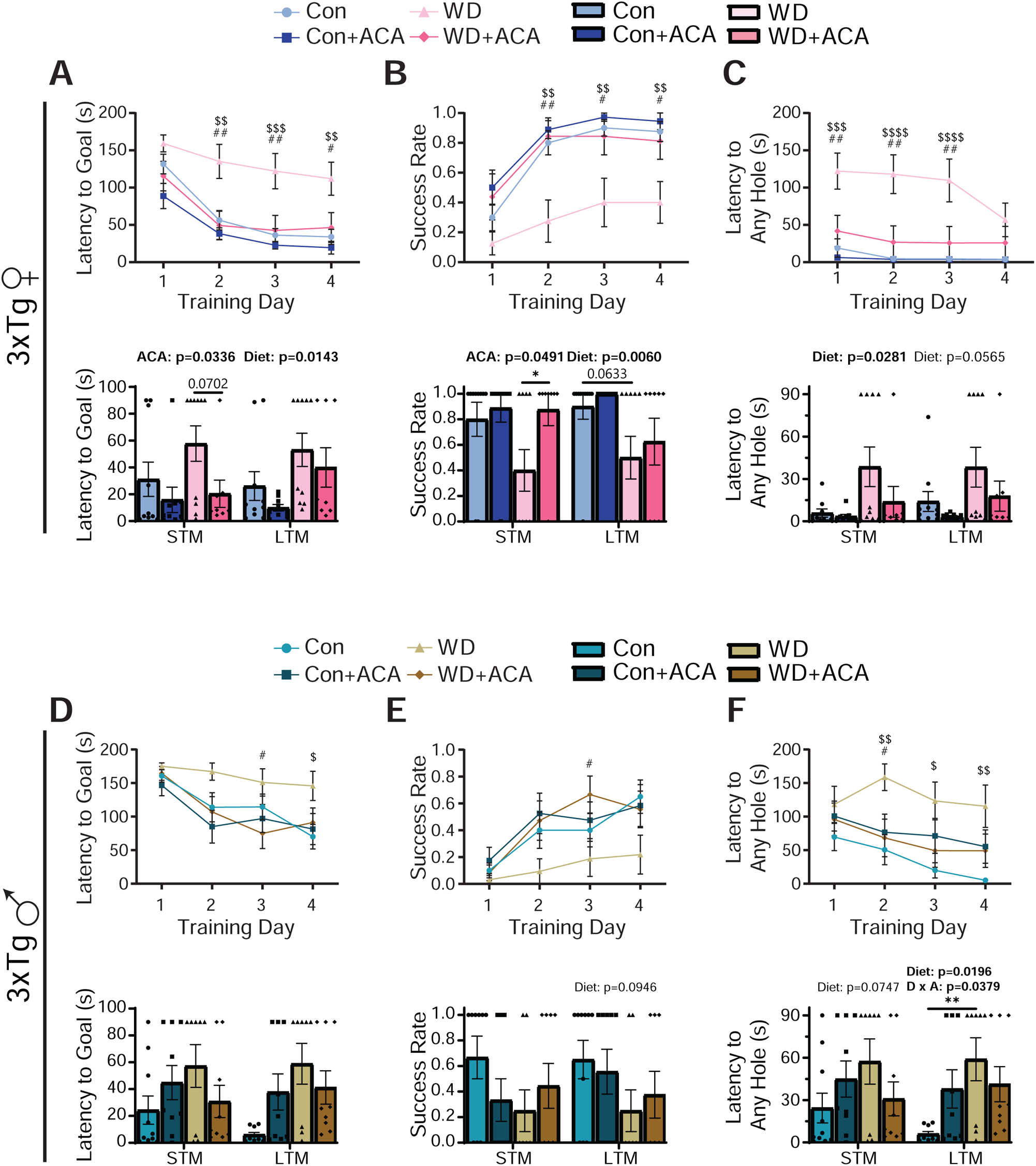
WD-induced impairments in spatial learning, memory, and anxiety in female and male 3xTg mice are reversed by acarbose. (A-F) Barnes maze was used to assess spatial learning and memory. (A, D) Latency to goal was measured in female (A) and male (D) mice. (B, E) Success rate of female (B) and male (E) mice. (C, F) Latency to any hole, a measure of anxiety, of female (C) and male (F) mice. (A-F) n = 8-10; statistics for the overall effects of diet, acarbose treatment (ACA), and the interaction represent the p-value from a 2-way ANOVA conducted for STM and LTM separately, with Sidak post-test, *p < 0.05 and **p < 0.01. For training days, statistics represent a pairwise 2-way ANOVA for the effects of diet group and training day with Sidak post-test, $p < 0.05 Con vs WD, $$p < 0.01 Con vs WD, $$$p < 0.001 Con vs WD, $$$$p < 0.0001 Con vs WD, #p < 0.05 WD vs WD+A, ##p < 0.01 WD vs WD+A. Data represented as mean ± SEM

Novel object recognition (NOR) was used to assess recognition memory (**Figs. S6A-D**). In female 3xTg mice, neither diet nor treatment had any effect on discrimination index (DI) or absolute DI although female 3xTg mice in general exhibited a stronger preference for the familiar object than the novel object, as supported by previous research showing that 3xTg mice exhibit neophobia (45) (**Fig. S6A**). In male 3xTg mice, WD significantly worsened discrimination index and neophobia in the long-term memory test, although those changes were not seen in the absolute DI (**Fig. S6B**). Overall, male 3xTg mice did not have as pronounced a neophobia as females. Female NTg mice did not exhibit any changes in DI or absolute DI, although they were much less neophobic than 3xTg females (**Fig. S6C**). NTg males did not exhibit any significant changes due to diet or acarbose alone, but rather showed an interaction effect wherein the combination of WD and acarbose was detrimental compared to WD-fed mice (**Fig. S6D**).

### WD impairs the survival of male 3xTg mice

Female 3xTg mice did not have any changes to survival with either WD or acarbose during the period of the experiment (**Fig. 9A**). Consistent with previous reports of our lab and others (46–49), we observed that male 3xTg mice had high levels of mortality during the period of the experiment, particularly when fed a WD (**Fig. 9B**). Male 3xTg mice had significantly higher mortality, with ending survival rates of 64%, 47%, 19%, and 30% in Con, Con+ACA, WD, and WD+ACA, respectively (**Fig. 9B**). WD was associated with double the mortality rate than Con in male mice. Acarbose did not improve survival in either Con-or WD-fed male 3xTg mice.

**Figure 9:**
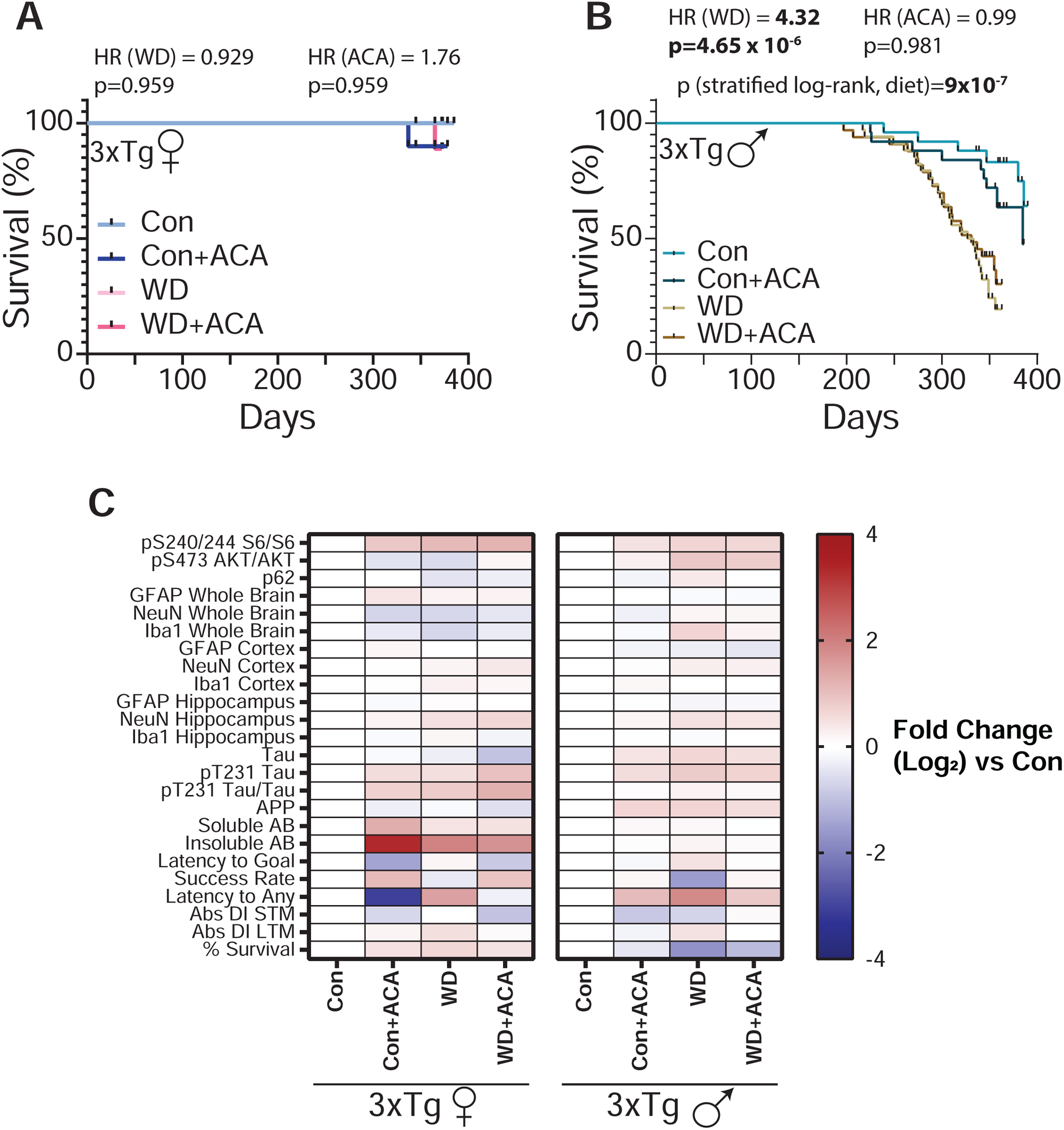
WD increases mortality in male 3xTg mice. (A-B) Kaplan-Meier plots of the survival of female (A) and male (B) 3xTg mice starting at 6 months of age. (A) n = 8-10 mice/group, (B) n = 8-34 mice/group. The overall effect of diet (WD) and acarbose (ACA) was determined using a Cox proportional hazards test (HR, hazard ratio). The two-tailed stratified log-rank *p*-value for the decrease in lifespan of 3xTg males as a result of WD is shown in (B). (C) Heat map representation of all the brain signaling, AD neuropathology, and cognitive parameters measured in female and male 3xTg mice; color represents the log_2_ fold-change vs female or male 3xTg mice fed control diet.

## Discussion

Metabolic health is an important risk factor for AD, and as such, alterations to diet that affect metabolic health may have a strong impact on AD pathology and cognitive function; dietary restrictions, for example, are beneficial for metabolic health and have also been shown in a variety of models to improve AD pathology and cognition (reviewed in (26)). WD has been proposed by many to worsen AD pathology, both on its own and through the development of obesity and T2DM (reviewed in (5)). WD feeding has been shown in a variety of mouse models to worsen AD pathology (5, 8, 9, 11). On the other hand, pharmaceutical interventions that are effective at treating either obesity or T2DM and promote improved glucose homeostasis may be beneficial for AD, suggesting that there may be untapped potential in already FDA approved pharmaceuticals.

Here we explore the use of the α-glucosidase inhibitor acarbose, which slows the uptake of carbohydrates in the small intestine and improves glycemic control in T2DM patients and is a geroprotective agent that extends the lifespan of mice, to slow or prevent the progression of AD in the 3xTg mouse model. We began acarbose treatment at 6 months of age, a point at which 3xTg mice in our animal colony have already begun to develop cognitive deficits and other aspects of AD pathology (49), in order to model a human beginning treatment after an early diagnosis. We administered acarbose to both NTg and 3xTg mice in the context of both Con and WD to examine the effects of acarbose in both wild-type mice and specifically in the context of AD with and without WD-induced metabolic syndrome.

As expected, we observed that WD worsened metabolic health in both NTg and 3xTg mice. WD significantly diminished spatial learning in the 3xTg mice, which was completely reversed by acarbose treatment in both sexes. Surprisingly, in contrast to other interventions that have been tested in mice, we saw that the effect of acarbose on brain mTOR signaling, autophagy, and AD pathology were fairly muted, highly dependent upon sex and strain, and did not clearly correlate with the benefits of acarbose on cognition of WD-fed mice.

WD increased body mass and more than doubled adiposity of all sexes and strains compared to their Con-fed counterparts. Acarbose reversed these effects, reducing body mass and adiposity to a level comparable to or even lower than Con-fed mice, despite acarbose-treated WD-fed mice eating more. These changes are likely due to changes in energy expenditure; WD decreased energy expenditure and acarbose restored energy expenditure to “normal” levels. With respect to glucose tolerance, WD-feeding impaired the glucose tolerance of female mice, an effect that was reversed by acarbose. While we did not observe an effect of WD on glucose tolerance in males, acarbose also improved the glucose tolerance of WD-fed males. Finally, sensitivity to IP insulin was not impacted by diet or treatment, except for an improvement of insulin sensitivity in Con-fed acarbose treated NTg male mice.

We assessed cognitive function using both Barnes maze and novel object recognition. Strikingly, WD strongly diminished spatial learning and memory and worsened anxiety in 3xTg mice of both sexes, and these effects were completely reversed by acarbose. In female mice, neither WD nor acarbose affected performance in the novel object recognition test, but in males WD worsened both object discrimination and neophobia in LTM. These results show a strong link between alterations to metabolic health and cognitive deficits.

To identify possible mechanisms between changes in metabolic health and cognition, we examined the effect of WD and acarbose on the mTOR nutrient signaling pathways through Western blotting of downstream targets. In 3xTg females, there was an overall effect of WD towards increased mTORC1 and decreased mTORC2 signaling in the brain, with acarbose reversing the effect on mTORC2. In male 3xTg mice, WD did not alter mTORC1 signaling but trended toward increased mTORC2 signaling. Acarbose treatment did not affect mTORC1 or mTORC2 signaling in male 3xTg mice. In the NTg mice, mTORC1 signaling is decreased in males fed a WD but not in females whereas mTORC2 signaling is increased in both sexes. These results suggest that changes in mTORC1 and mTORC2 activity in the brain in response to WD and acarbose are strongly dependent on sex and strain and are likely not a strong driver of the changes in cognition.

p62 is an autophagy receptor and is a key player in the clearance of neurofibrillary tangles through direct binding and preparation for degradation (50–52). Increased p62 levels typically indicate inhibition of autophagy due to retention of p62 in the cell as opposed to degradation in a lysosome (52, 53). We found clear sex- and strain-dependent differences in the autophagic response to WD and acarbose. Male 3xTg mice fed a WD had increased levels of p62 (p=0.1025), suggesting an impairment in autophagy, that was ameliorated (p=0.0906) by acarbose. These results correlated with changes in cognition. 3xTg female mice, however, exhibit increased (p=0.0613) autophagy during WD feeding which coincides with the decrease in S473 AKT phosphorylation seen in WD-fed mice. AKT has been shown to be a potent mTORC1-independent inhibitor of autophagy (29) so decreased S473 AKT phosphorylation via WD feeding may be responsible for the decreased p62 levels seen in females fed a WD. However, the changes in autophagy in females do not correlate with the cognition results as WD worsened cognition despite an increase in autophagy. In NTg mice, neither WD nor acarbose altered p62 levels.

Next, we assessed the impact of WD and acarbose on aspects of AD pathology in the 3xTg mice. We measured the levels of different brain cell types (astrocytes, neurons, and microglia) in both the whole brain and specifically in the cortex and hippocampus and measured both tau and amyloid pathology in the whole brain. Once again, WD and acarbose exhibited dramatic sex-dependent differences. GFAP (astrocyte marker) levels in the cortex and hippocampus are unchanged in female mice, but in male mice are decreased by WD feeding, which may suggest impairments in glucose utilization in the brain (54). 3xTg mice have been shown to have increased Iba1 (microglia) levels compared to NTg controls (34) and Iba1 levels are increased in WD-fed male mice, indicating increased inflammation which is commonly associated with AD (34, 55, 56) and may contribute to neuronal dysfunction. Acarbose reversed the WD-induced increase in Iba1 in male mice. In addition, male mice exhibited a morphological change to their microglial populations in the hippocampus, with WD lending to a more reactive state which is partially reversed by acarbose. In concordance with the increased autophagy seen in WD-fed female 3xTg mice, they also exhibited a non-significant (p=0.0568) overall trend of WD to decrease Aβ40 levels, the clearance of which autophagy is primarily responsible. In addition, the proportion of T231 tau phosphorylation is increased in WD-fed female mice. In male 3xTg mice, both tau and T231 tau phosphorylation are increased in WD-fed mice, as well as insoluble Aβ40 levels. While not increased by WD, soluble Aβ40 levels are decreased by acarbose treatment. As a whole, female mice did not show a strong correlation between AD pathology and cognition, exhibiting worsened cognition on WD despite improvements in autophagy, Iba1, and Aβ40 levels. Males showed a stronger link between AD pathology and cognition via autophagy, Iba1, tau, and Aβ40 levels.

These results suggest that in 3xTg males, WD leads to impaired metabolic health, autophagy, increased inflammation, and increased Aβ levels which lead to worsened cognition, all of which are ameliorated by acarbose treatment. In females, the relationship between WD, acarbose, and AD pathology is more complex. WD did indeed worsen metabolic health and cognitive deficits which were completely reversed by acarbose treatment, however, female mice on WD exhibit an increase in autophagy and corresponding decreases in both inflammation and Aβ levels. Increased autophagy in 3xTg females could be a compensatory mechanisms to offset AD pathology; studies in APP/PS1 mice have found that early in amyloid pathology, autophagy is increased in an attempt to offset initial Aβ deposition (57, 58). As seen previously in our lab, Aβ pathology is still in its nascency at 12-13 months of age, and true Aβ aggregation doesn’t occur until closer to 15 months of age (49), which supports the idea of increased autophagy in the female WD-fed mice as a compensatory mechanism. Female mice exhibited similar levels of soluble amyloid as males but had approximately 100 times greater levels of insoluble amyloid, suggesting greater amyloid pathology. This is reflective of amyloid pathology in humans, wherein females tend to see more extreme AD pathology than males (59). Interestingly, previous research on acarbose have shown that acarbose has more dramatic improvements on lifespan and metabolic health in males than females, albeit body and fat mass was improved to the same extent or more in some female mice (16, 17, 27, 32, 60). In both the 3xTg and NTg mice, acarbose was equally effective at improving body mass and adiposity, energy expenditure, and glycemic control in males and females. Cognition was actually improved more dramatically in females than males. These results highlight important sex differences in AD pathology and response to both diet and pharmaceuticals.

Further research will be needed to determine the actual mechanism responsible for the alterations to cognition, which are overall more strongly correlated with changes to metabolic health than AD pathology. One possible mechanism is the improvement of glucose homeostasis. As shown here, acarbose improves glucose tolerance in mice fed WD. Glucose homeostasis and glucose uptake in the brain is perturbed during AD, so improvements to peripheral glucose signaling may also coincide with improvements in brain glucose signaling as well (3). In addition, acarbose has been previously shown to improve non-alcoholic steatohepatitis (NASH) in rodent models (61, 62). Recently, it has been reported that liver dysfunction, and in particular NASH, contribute to worsened AD pathology in mice (63, 64), and thus it is possible that the beneficial effects of acarbose on cognition we observed are due in part to improved liver health.

Limitations of the current study include the use of 3xTg mice as the AD model. Although 3xTg mice are a valuable tool for testing the impact of diet and interventions on AD pathology and develop both tau and Aβ pathology, the model does have its limitations. First, like most rodent models of AD, 3xTg are a model of early onset, or familial, AD. This limits the translatability of work in 3xTg mice to late-onset AD patients, although there is still the potential for 3xTg mice to be used to study how diet and interventions may impact AD pathology directly. In addition, 3xTg mice lack a robust aggregation of Aβ plaques at 12-13 months of age, which limits our ability to determine how WD or acarbose impact Aβ pathology. Aging the mice longer as we have done in other studies was difficult due to the exceptionally high mortality of WD-fed 3xTg male mice. Utilizing additional AD mouse models may give additional insight into how WD or acarbose may affect tau or Aβ pathology directly. Finally, we utilized only a single mixed genetic background, and it has become clear that strain is a critical factor in the response to both diets and drugs (65–68); it is possible that the effect of WD and acarbose on AD pathology and on cognition may differ depending on genetic background.

In conclusion, we have shown that WD has negative effects on metabolic health and cognition in 3xTg mice, which are reversed by acarbose treatment. We also found that the effects of WD and acarbose on molecular signaling in the brain and on AD pathology do not correlate clearly with the effects on cognition and are strongly sex dependent. Our results suggest that the benefits of acarbose on cognition of WD-fed 3xTg mice are mediated not by direct action on AD pathology, but instead by improvements in metabolic health. Our results support the idea that already FDA approved pharmaceuticals for other age-related diseases may be beneficial for the prevention or treatment of AD pathology — particularly the use of anti-diabetes drugs such as acarbose.

## Materials and Methods

### Animals

All procedures were performed in accordance with institutional guidelines and were approved by the Institutional Animal Care and Use Committee (IACUC) of the William S. Middleton Memorial Veterans Hospital (Madison, WI, USA) and the University of Wisconsin-Madison IACUC (Madison, WI, USA). Male and female homozygous 3xTg-AD and B6129SF2/J mice were originally obtained from The Jackson Laboratory/Mutant Mouse Resource & Research Centers and were bred and maintained with food and water available *ad libitum*. All mice were maintained at a temperature of approximately 22°C, with a 12:12 light/dark cycle. Health checks were completed on all mice daily. Mice were acclimatized to our experimental facility for one week before experiment start and were housed 2-3 per cage. At 2 months of age, half of the mice of each genotype and sex were randomly assigned to receive either Control diet (Con; Envigo Global 2018) or Western diet (WD; TD.88137). At 6 months of age, the mice were then split further into different groups: half of the Con-fed mice remained on Con while the other half received a custom diet consisting of Teklad 2018 plus 1000ppm of acarbose (Con+ACA; TD.190479). Half of the WD-fed mice remained on WD awhile the other half received a custom diet consisting of the WD plus acarbose (WD+ACA; TD.210479). All diets were obtained from Inotivo (formerly Envigo). Full diet descriptions, compositions, and item numbers are provided in **Table S1**. Randomization of mice to groups was performed at cage level to ensure all groups had approximately the same starting weight and body composition.

### In vivo procedures

Glucose tolerance tests were performed by fasting the mice overnight for 16 hours then injecting glucose (1 g kg^−1^) intraperitoneally (i.p.) (69, 70). Insulin tolerance tests were performed by fasting the mice for 4 hours then injecting insulin I.P. (0.75 U kg^−1^). Blood glucose measurements were taking using a Bayer Contour blood glucose meter (Bayer, Leverkusen, Germany) and test strips. Mouse body composition was analyzed at several points throughout the study using an EchoMRI Body Composition Analyzer (EchoMRI, Houston, TX, USA). Metabolic parameters (O_2_, CO_2_, food composition, respiratory exchange ratio (RER), and energy expenditure) and activity tracking were analyzed using a Columbus Instruments Oxymax/CLAMS metabolic chamber system. Mice were acclimated to the chamber cages for ∼24 hours and data from a subsequent, continuous 24 hour period was recorded and analyzed. Food consumption in home cages was measured by moving mice to clean cages, filling the hopper with a measured quantity of fresh diet, and measuring the remainder 3-4 days later. The amount of food consumed was adjusted for the number of mice in each cage, the number of days that had passed, and the relative weights of the mice. Mice were euthanized by cervical dislocation after a 3 hour fast and tissues for molecular analysis were flash-frozen in liquid nitrogen or fixed and prepared as described below. For the brain, the right hemisphere was fixed in formalin for histology whereas the left hemisphere was flash-frozen for Western blot and ELISA analyses.

### Behavioral Assays

All mice underwent behavioral phenotyping at 11-12 months of age. All behavioral assays were recorded and analyzed utilizing the EthoVision XT animal tracking software by Noldus. Before each procedure the mice were allowed to acclimate to the behavioral testing room for 30 minutes. The arenas and objects used were all cleaned with 70% ethanol to remove olfactory cues between mice and runs.

For Novel Object Recognition (NOR), mice were first habituated to the empty arena with no objects. Approximately 24 hours later, an acquisition trial was performed where the mice were placed in the arena with two of the same object, equidistant from the mouse, hereafter referred to as “Object A”. The mouse was allowed to explore the two objects for 5 minutes before returning to the home cage. One hour after the acquisition trial, a short-term memory (STM) test was performed during which the mouse was placed into the arena with one of object A as well as a single novel object, or “Object B”, and allowed to explore for 5 minutes. The following day, 24 hours after the STM test, a long-term memory (LTM) test was performed. During the LTM test, the mouse was placed in the arena with one of object A and a single novel object, “Object C”, and allowed to explore for 5 minutes. Results were quantified by calculating discrimination index (DI) by dividing the time spent exploring the novel object by the time spent examining either object. A DI value between −0.2 and 0.2 indicates no difference in time spent examining each object. A positive value indicates greater time was spent examining the novel object whereas a negative value indicates greater time was spent examining the familiar object. Due to the nature of the DI calculation, absolute DI is also calculated to separate discrimination from object preference.

For Barnes maze, the test involved 4 phases: habituation, acquisition, and STM and LTM memory tests. During habituation, the mice were placed on the maze and immediately led to the escape box where they remained for 2 minutes before returning to their home cage. After approximately 15 minutes, the mice underwent their first acquisition trial in which they were allowed to explore the maze until either they found the escape box or 3 minutes had elapsed. If the mouse did not find the escape box in 3 minutes, they were gently led to the escape box. Regardless of how the mouse reached the escape box, they remained there for 1 minute after every trial. Over the first four days, four acquisition trials were performed each day with an inter-trial interval of 15 minutes between each trial. On days 5 (STM test) and 12 (LTM test), the escape box was removed and the mouse was placed on the maze for 90s to determine how well it was able to recall where the escape box was. The latency to goal, success rate, and latency to any hole were analyzed. For success rate, mice were determined to be successful if they made it to the goal in 90 seconds or less and as such were scored a 1. Mice that did not make it to the goal within 90 seconds were scored a 0.

### Immunoblotting

Tissue samples (brain and liver) were lysed in cold RIPA buffer supplemented with phosphatase inhibitor and protease inhibitor cocktail tablets (Thermo Fisher Scientific, Waltham, MA, USA) using a FastPrep 24 (M.P. Biomedicals, Santa Ana, CA, USA) with bead-beating tubes from (VWR, Radnor, PA, USA) and zirconium ceramic oxide beads from (Thermo Fisher Scientific, Waltham, MA, USA). Protein lysates were then centrifuged at 13,300 rpm for 10 min and the supernatant was collected. Protein concentration was determined by Bradford Assay (Pierce Biotechnology, Waltham, MA, USA). 20-60 μg protein was separated on 8%, 10%, or 16% tris-glycine gels (ThermoFisher Scientific, Waltham, MA, USA) and transferred to PVDF membrane (EMD Millipore, Burlington, MA, USA). The phosphorylation of mTOR substrates including pS240/244 S6 and pS473 AKT were assessed in the brain and liver along with p62 protein receptor and markers for brain cell populations, specifically GFAP, NeuN, and Iba1. Markers for AD pathology were also assessed by Western blot, specifically targeting p-Thr231 tau, total tau, and APP. Antibody vendors, catalog numbers and the dilution used is provided in **Table S2**. Imaging was performed using a Bio-Rad Chemidoc MP imaging station (Bio-Rad, Hercules, CA, USA). Quantification was performed by densitometry using NIH ImageJ software.

### Immunohistochemistry for AD neuropathology markers

Formalin-fixed brains were paraffin embedded and cut into 5 μm sections which were affixed to Superfrost Plus microscope slides (ThermoFisher Scientific, Waltham, MA, USA). Sections were deparaffinized and rehydrated then stained with the following primary antibodies: anti-GFAP (ThermoFisher; #PIMA512023; 1:1,000), anti-Iba1 (Abcam; #ab178847; 1:1,000), and anti-NeuN. Sections were imaged using an EVOS microscope (ThermoFisher Scientific, Waltham, MA, USA) at magnifications of 10X and 20X. Image-J was used for quantification by converting images into binary images via an intensity threshold and positive area was quantified.

### ELISA

Amyloid-β 40 (Aβ40) in the brain was quantified using a human Aβ40 ELISA kit (Invitrogen, USA, #KHB3482) in accordance with manufacturer’s instructions.

### Statistical Analyses

All statistical analyses were conducted using Prism, version 9 and 10 (GraphPad Software Inc., San Diego, CA, USA). Tests involving multiple factors were analyzed by either a two-way analysis of variance (ANOVA) with diet and genotype as variables or by one-way ANOVA, followed by a Dunnett’s, Tukey-Kramer, or Sidak’s post-hoc test as specified in the figure legends.

Survival analyses were conducted in R using the “survival” package (Therneau, 2015). Kaplan– Meir survival analysis of 3xTg mice was performed with log-rank comparisons stratified by sex and diet. Cox proportional hazards analysis of 3xTg mice was performed using sex and diet as covariates. Alpha was set at 5% (p < .05 considered to be significant). Data are presented as the mean ± SEM unless otherwise specified.

## Acknowledgements

We thank all members of the Lamming lab for their feedback. The Lamming lab is supported in part by the NIA (AG056771, AG062328, AG081482, and AG084156), the NIDDK (DK125859), by a grant from the Alzheimer’s Association (23AARG-1029665), and by startup funds from UW-Madison. RB was supported in part by F31AG081115. MMS was supported in part by a Supplement to Promote Diversity in Health-Related Research RF1AG056771-06S1. MET was supported in part by a Supplement to Promote Diversity in Health-Related Research R01AG062328-03S1 and by F32AG077916. CLG was supported in part by Dalio Philanthropies, a Glenn Foundation for Medical Research Postdoctoral Fellowship, and by grant HF-AGE AGE-009 from the Hevolution Foundation to CLG. MFC was supported in part by F31AG082504. C-YY was supported in part by a training grant from the NIA (T32 AG000213) and by F32AG077916. HHP was supported in part by F31AG066311. The Puglielli lab is supported in part by the NINDS (NS094154), the NIGMS (GM148487) and the NIA (AG078794). The Lamming lab was supported in part by the U.S. Department of Veterans Affairs (I01-BX004031 and IS1-BX005524), and this work was supported using facilities and resources from the William S. Middleton Memorial Veterans Hospital. The content is solely the responsibility of the authors and does not necessarily represent the official views of the NIH. This work does not represent the views of the Department of Veterans Affairs or the United States Government.

## Author Contributions

MMS, RB, MJ, SC, MC, C-YY, IG, YL, DV, MFC, MET, CLG, and MJR conducted the experiments. MMS, RB, DV, and DWL analyzed the data. MMS, RB, DV, LP, and DWL wrote and edited the manuscript.

## References

1. Association As. Alzheimer’s disease facts and figures. Alzheimers Dement. 2023;19(4):1598–695.

2. Sibener L, Zaganjor I, Snyder HM, Bain LJ, Egge R, Carrillo MC. Alzheimer’s Disease prevalence, costs, and prevention for military personnel and veterans. Alzheimer’s & dementia : the journal of the Alzheimer’s Association. 2014;10(3 Suppl):S105–10.

3. Duran-Aniotz C, Hetz C. Glucose Metabolism: A Sweet Relief of Alzheimer’s Disease. Curr Biol. 2016;26(17):R806–9.

4. Rakhra V, Galappaththy SL, Bulchandani S, Cabandugama PK. Obesity and the Western Diet-How We Got Here. Mo Med. 2020;117(6):536--8.

5. Wieckowska-Gacek A, Mietelska-Porowska A, Wydrych M, Wojda U. Western diet as a trigger of Alzheimer’s disease: From metabolic syndrome and systemic inflammation to neuroinflammation and neurodegeneration. Ageing Res Rev. 2021;70:101397.

6. Kopp W. How Western Diet And Lifestyle Drive The Pandemic Of Obesity And Civilization Diseases. Diabetes Metab Syndr Obes. 2019;12:2221–36.

7. Demaria TM, Crepaldi LD, Costa-Bartuli E, Branco JR, Zancan P, Sola-Penna M. Once a week consumption of Western diet over twelve weeks promotes sustained insulin resistance and non-alcoholic fat liver disease in C57BL/6 J mice. Sci Rep. 2023;13(1):3058.

8. Graham LC, Harder JM, Soto I, de Vries WN, John SW, Howell GR. Chronic consumption of a western diet induces robust glial activation in aging mice and in a mouse model of Alzheimer’s disease. Sci Rep. 2016;6:21568.

9. Kanoski SE, Davidson TL. Western diet consumption and cognitive impairment: links to hippocampal dysfunction and obesity. Physiol Behav. 2011;103(1):59–68.

10. Sullivan PM. Influence of Western diet and APOE genotype on Alzheimer’s disease risk. Neurobiol Dis. 2020;138:104790.

11. Wieckowska-Gacek A, Mietelska-Porowska A, Chutoranski D, Wydrych M, Dlugosz J, Wojda U. Western Diet Induces Impairment of Liver-Brain Axis Accelerating Neuroinflammation and Amyloid Pathology in Alzheimer’s Disease. Front Aging Neurosci. 2021;13:654509.

12. Altay M. Acarbose is again on the stage. World J Diabetes. 2022;13(1):1–4.

13. Wehmeier UF, Piepersberg W. Biotechnology and molecular biology of the alpha-glucosidase inhibitor acarbose. Appl Microbiol Biotechnol. 2004;63(6):613–25.

14. Chiasson JL, Josse RG, Gomis R, Hanefeld M, Karasik A, Laakso M, et al. Acarbose for prevention of type 2 diabetes mellitus: the STOP-NIDDM randomised trial. Lancet. 2002;359(9323):2072-7.

15. Dodds SG, Parihar M, Javors M, Nie J, Musi N, Dave Sharp Z, et al. Acarbose improved survival for Apc(+/Min) mice. Aging Cell. 2020;19(2):e13088.

16. Harrison DE, Strong R, Alavez S, Astle CM, DiGiovanni J, Fernandez E, et al. Acarbose improves health and lifespan in aging HET3 mice. Aging Cell. 2019;18(2):e12898.

17. Harrison DE, Strong R, Allison DB, Ames BN, Astle CM, Atamna H, et al. Acarbose, 17-alpha-estradiol, and nordihydroguaiaretic acid extend mouse lifespan preferentially in males. Aging Cell. 2014;13(2):273–82.

18. Wang Z, Wang J, Hu J, Chen Y, Dong B, Wang Y. A comparative study of acarbose, vildagliptin and saxagliptin intended for better efficacy and safety on type 2 diabetes mellitus treatment. Life Sci. 2021;274:119069.

19. Shen Z, Hinson A, Miller RA, Garcia GG. Cap-independent translation: A shared mechanism for lifespan extension by rapamycin, acarbose, and 17alpha-estradiol. Aging Cell. 2021;20(5):e13345.

20. Simcox J, Lamming DW. The central moTOR of metabolism. Dev Cell. 2022;57(6):691–706.

21. Mannick JB, Lamming DW. Targeting the biology of aging with mTOR inhibitors. Nat Aging. 2023;3(6):642–60.

22. Caccamo A, Branca C, Talboom JS, Shaw DM, Turner D, Ma L, et al. Reducing Ribosomal Protein S6 Kinase 1 Expression Improves Spatial Memory and Synaptic Plasticity in a Mouse Model of Alzheimer’s Disease. J Neurosci. 2015;35(41):14042–56.

23. Caccamo A, Magri A, Medina DX, Wisely EV, Lopez-Aranda MF, Silva AJ, et al. mTOR regulates tau phosphorylation and degradation: implications for Alzheimer’s disease and other tauopathies. Aging Cell. 2013;12(3):370–80.

24. Caccamo A, Majumder S, Richardson A, Strong R, Oddo S. Molecular interplay between mammalian target of rapamycin (mTOR), amyloid-beta, and Tau: effects on cognitive impairments. J Biol Chem. 2010;285(17):13107–20.

25. Spilman P, Podlutskaya N, Hart MJ, Debnath J, Gorostiza O, Bredesen D, et al. Inhibition of mTOR by rapamycin abolishes cognitive deficits and reduces amyloid-beta levels in a mouse model of Alzheimer’s disease. PLoS One. 2010;5(4):e9979.

26. Sonsalla MM, Lamming DW. Geroprotective interventions in the 3xTg mouse model of Alzheimer’s disease. Geroscience. 2023;45(3):1343–81.

27. Garratt M, Bower B, Garcia GG, Miller RA. Sex differences in lifespan extension with acarbose and 17-alpha estradiol: gonadal hormones underlie male-specific improvements in glucose tolerance and mTORC2 signaling. Aging Cell. 2017;16(6):1256–66.

28. Querfurth H, Lee HK. Mammalian/mechanistic target of rapamycin (mTOR) complexes in neurodegeneration. Mol Neurodegener. 2021;16(1):44.

29. Ballesteros-Alvarez J, Andersen JK. mTORC2: The other mTOR in autophagy regulation. Aging Cell. 2021;20(8):e13431.

30. Markaki M, Tavernarakis N. Metabolic control by target of rapamycin and autophagy during ageing - a mini-review. Gerontology. 2013;59(4):340–8.

31. Meijer AJ, Lorin S, Blommaart EF, Codogno P. Regulation of autophagy by amino acids and MTOR-dependent signal transduction. Amino Acids. 2015;47(10):2037–63.

32. Wink L, Miller RA, Garcia GG. Rapamycin, Acarbose and 17alpha-estradiol share common mechanisms regulating the MAPK pathways involved in intracellular signaling and inflammation. Immun Ageing. 2022;19(1):8.

33. Sadagurski M, Cady G, Miller RA. Anti-aging drugs reduce hypothalamic inflammation in a sex-specific manner. Aging Cell. 2017;16(4):652–60.

34. Belfiore R, Rodin A, Ferreira E, Velazquez R, Branca C, Caccamo A, et al. Temporal and regional progression of Alzheimer’s disease-like pathology in 3xTg-AD mice. Aging Cell. 2019;18(1):e12873.

35. Oddo S, Caccamo A, Kitazawa M, Tseng BP, LaFerla FM. Amyloid deposition precedes tangle formation in a triple transgenic model of Alzheimer’s disease. Neurobiol Aging. 2003;24(8):1063–70.

36. Oddo S, Caccamo A, Shepherd JD, Murphy MP, Golde TE, Kayed R, et al. Triple-Transgenic Model of Alzheimer’s Disease with Plaques and Tangles. Neuron. 2003;39(3):409–21.

37. Barron AM, Rosario ER, Elteriefi R, Pike CJ. Sex-specific effects of high fat diet on indices of metabolic syndrome in 3xTg-AD mice: implications for Alzheimer’s disease. PLoS One. 2013;8(10):e78554.

38. Julien C, Tremblay C, Phivilay A, Berthiaume L, Emond V, Julien P, et al. High-fat diet aggravates amyloid-beta and tau pathologies in the 3xTg-AD mouse model. Neurobiol Aging. 2010;31(9):1516–31.

39. Robison LS, Gannon OJ, Thomas MA, Salinero AE, Abi-Ghanem C, Poitelon Y, et al. Role of sex and high-fat diet in metabolic and hypothalamic disturbances in the 3xTg-AD mouse model of Alzheimer’s disease. J Neuroinflammation. 2020;17(1):285.

40. Herrera JJ, Pifer K, Louzon S, Leander D, Fiehn O, Day SM, et al. Early or Late-Life Treatment With Acarbose or Rapamycin Improves Physical Performance and Affects Cardiac Structure in Aging Mice. J Gerontol A Biol Sci Med Sci. 2023;78(3):397–406.

41. Hasek BE, Stewart LK, Henagan TM, Boudreau A, Lenard NR, Black C, et al. Dietary methionine restriction enhances metabolic flexibility and increases uncoupled respiration in both fed and fasted states. Am J Physiol Regul Integr Comp Physiol. 2010;299(3):R728–39.

42. Pak HH, Haws SA, Green CL, Koller M, Lavarias MT, Richardson NE, et al. Fasting drives the metabolic, molecular and geroprotective effects of a calorie-restricted diet in mice. Nat Metab. 2021;3(10):1327–41.

43. Sun YX, Ji X, Mao X, Xie L, Jia J, Galvan V, et al. Differential activation of mTOR complex 1 signaling in human brain with mild to severe Alzheimer’s disease. Journal of Alzheimer’s disease : JAD. 2014;38(2):437–44.

44. Perluigi M, Di Domenico F, Barone E, Butterfield DA. mTOR in Alzheimer disease and its earlier stages: Links to oxidative damage in the progression of this dementing disorder. Free Radic Biol Med. 2021;169:382–96.

45. Li Y, Zheng W, Lu Y, Zheng Y, Pan L, Wu X, et al. BNIP3L/NIX-mediated mitophagy: molecular mechanisms and implications for human disease. Cell Death Dis. 2021;13(1):14.

46. Gimenez-Llort L, Arranz L, Mate I, De la Fuente M. Gender-specific neuroimmunoendocrine aging in a triple-transgenic 3xTg-AD mouse model for Alzheimer’s disease and its relation with longevity. Neuroimmunomodulation. 2008;15(4-6):331–43.

47. Garcia-Mesa Y, Colie S, Corpas R, Cristofol R, Comellas F, Nebreda AR, et al. Oxidative Stress Is a Central Target for Physical Exercise Neuroprotection Against Pathological Brain Aging. J Gerontol A Biol Sci Med Sci. 2016;71(1):40–9.

48. Hirata-Fukae C, Li HF, Hoe HS, Gray AJ, Minami SS, Hamada K, et al. Females exhibit more extensive amyloid, but not tau, pathology in an Alzheimer transgenic model. Brain research. 2008;1216:92–103.

49. Babygirija R, Sonsalla MM, Mill J, James I, Han JH, Green CL, et al. Protein restriction slows the development and progression of pathology in a mouse model of Alzheimer’s disease. Nature communications. 2024;15(1):5217.

50. Ramesh Babu J, Lamar Seibenhener M, Peng J, Strom AL, Kemppainen R, Cox N, et al. Genetic inactivation of p62 leads to accumulation of hyperphosphorylated tau and neurodegeneration. J Neurochem. 2008;106(1):107–20.

51. Salminen A, Kaarniranta K, Haapasalo A, Hiltunen M, Soininen H, Alafuzoff I. Emerging role of p62/sequestosome-1 in the pathogenesis of Alzheimer’s disease. Prog Neurobiol. 2012;96(1):87–95.

52. Su H, Wang X. Autophagy and p62 in cardiac protein quality control. Autophagy. 2011;7(11):1382–3.

53. Bjorkoy G, Lamark T, Pankiv S, Overvatn A, Brech A, Johansen T. Monitoring autophagic degradation of p62/SQSTM1. Methods Enzymol. 2009;452:181–97.

54. Machado LS, da Rocha AS, Soares C, de Souza DG, Ramos VG, Bellaver B, et al. GFAP expression levels are associated with brain glucose metabolism in a rat model of Alzheimer’s disease. Alzheimer’s & Dementia. 2022;18(S5).

55. Heneka MT, Carson MJ, El Khoury J, Landreth GE, Brosseron F, Feinstein DL, et al. Neuroinflammation in Alzheimer’s disease. Lancet Neurol. 2015;14(4):388–405.

56. Janelsins MC, Mastrangelo MA, Oddo S, LaFerla FM, Federoff HJ, Bowers WJ. Early correlation of microglial activation with enhanced tumor necrosis factor-alpha and monocyte chemoattractant protein-1 expression specifically within the entorhinal cortex of triple transgenic Alzheimer’s disease mice. J Neuroinflammation. 2005;2:23.

57. Yu WH, Cuervo AM, Kumar A, Peterhoff CM, Schmidt SD, Lee JH, et al. Macroautophagy--a novel Beta-amyloid peptide-generating pathway activated in Alzheimer’s disease. J Cell Biol. 2005;171(1):87–98.

58. de la Cueva M, Antequera D, Ordonez-Gutierrez L, Wandosell F, Camins A, Carro E, et al. Amyloid-beta impairs mitochondrial dynamics and autophagy in Alzheimer’s disease experimental models. Sci Rep. 2022;12(1):10092.

59. Oveisgharan S, Arvanitakis Z, Yu L, Farfel J, Schneider JA, Bennett DA. Sex differences in Alzheimer’s disease and common neuropathologies of aging. Acta Neuropathol. 2018;136(6):887–900.

60. Strong R, Miller RA, Antebi A, Astle CM, Bogue M, Denzel MS, et al. Longer lifespan in male mice treated with a weakly estrogenic agonist, an antioxidant, an alpha-glucosidase inhibitor or a Nrf2-inducer. Aging Cell. 2016;15(5):872–84.

61. Lieber CS, Leo MA, Mak KM, Xu Y, Cao Q, Ren C, et al. Acarbose attenuates experimental non-alcoholic steatohepatitis. Biochem Biophys Res Commun. 2004;315(3):699–703.

62. Xing Y, Ren X, Li X, Sui L, Shi X, Sun Y, et al. Baicalein Enhances the Effect of Acarbose on the Improvement of Nonalcoholic Fatty Liver Disease Associated with Prediabetes via the Inhibition of De Novo Lipogenesis. J Agric Food Chem. 2021;69(34):9822–36.

63. Kim DG, Krenz A, Toussaint LE, Maurer KJ, Robinson SA, Yan A, et al. Non-alcoholic fatty liver disease induces signs of Alzheimer’s disease (AD) in wild-type mice and accelerates pathological signs of AD in an AD model. J Neuroinflammation. 2016;13:1.

64. Estrada LD, Ahumada P, Cabrera D, Arab JP. Liver Dysfunction as a Novel Player in Alzheimer’s Progression: Looking Outside the Brain. Front Aging Neurosci. 2019;11:174.

65. Green CL, Lamming DW. We are more than what we eat. Nat Metab. 2021;3(9):1144–5.

66. Green CL, Pak HH, Richardson NE, Flores V, Yu D, Tomasiewicz JL, et al. Sex and genetic background define the metabolic, physiologic, and molecular response to protein restriction. Cell Metab. 2022;34(2):209–26 e5.

67. Barrington WT, Wulfridge P, Wells AE, Rojas CM, Howe SYF, Perry A, et al. Improving Metabolic Health Through Precision Dietetics in Mice. Genetics. 2018;208(1):399–417.

68. Roy S, Sleiman MB, Jha P, Ingels JF, Chapman CJ, McCarty MS, et al. Gene-by-environment modulation of lifespan and weight gain in the murine BXD family. Nat Metab. 2021;3(9):1217–27.

69. Bellantuono I, de Cabo R, Ehninger D, Di Germanio C, Lawrie A, Miller J, et al. A toolbox for the longitudinal assessment of healthspan in aging mice. Nature protocols. 2020;15(2):540–74.

70. Yu D, Yang SE, Miller BR, Wisinski JA, Sherman DS, Brinkman JA, et al. Short-term methionine deprivation improves metabolic health via sexually dimorphic, mTORC1-independent mechanisms. FASEB J. 2018;32(6):3471–82.

