## Supplemental Figures and Table Legends for "Acarbose ameliorates Western diet-induced metabolic and cognitive impairments in the 3xTg mouse model of Alzheimer’s disease"

### Figure S1

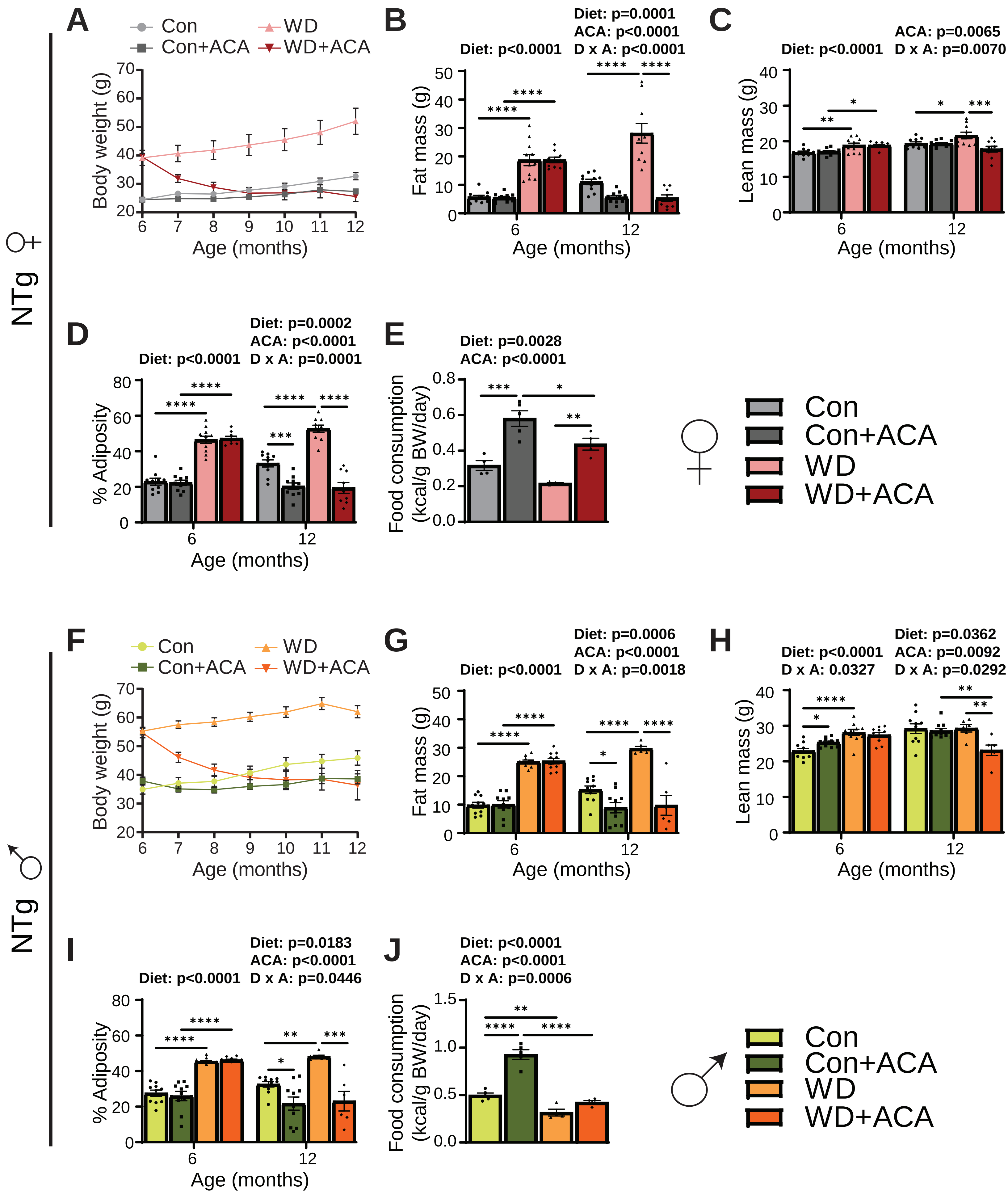

NTg ♀
A
Con
Con+ACA
WD
WD+ACA
Body weight (g)
Age (months)
B
Diet: p<0.0001
Diet: p=0.0001
ACA: p<0.0001
D x A: p<0.0001
Fat mass (g)
Age (months)
C
Diet: p<0.0001
ACA: p=0.0065
D x A: p=0.0070
Lean mass (g)
Age (months)
D
Diet: p<0.0001
Diet: p=0.0002
ACA: p<0.0001
D x A: p=0.0001
% Adiposity
Age (months)
E
Diet: p=0.0028
ACA: p<0.0001
Food consumption (kcal/g BW/day)
Con
Con+ACA
WD
WD+ACA
NTg ♂
F
Con
Con+ACA
WD
WD+ACA
Body weight (g)
Age (months)
G
Diet: p<0.0001
Diet: p=0.0006
ACA: p<0.0001
D x A: p=0.0018
Fat mass (g)
Age (months)
H
Diet: p<0.0001
D x A: 0.0327
Diet: p=0.0362
ACA: p=0.0092
D x A: p=0.0292
Lean mass (g)
Age (months)
I
Diet: p<0.0001
Diet: p=0.0183
ACA: p<0.0001
D x A: p=0.0446
% Adiposity
Age (months)
J
Diet: p<0.0001
ACA: p<0.0001
D x A: p=0.0006
Food consumption (kcal/g BW/day)
Con
Con+ACA
WD
WD+ACA

**Supplementary Figure 1: WD-induced changes in the body mass and adiposity of NTg mice are ameliorated by acarbose.**

(A-E) The body weight (A) of female NTg mice was measured over the course of the experiment while fat (B) and lean mass (C) were determined at both the beginning and end of the experiment using EchoMRI and adiposity (D) was calculated; n = 6-10 mice/group. (E) Food consumption was measured in home cages at 10 months of age and normalized to body weight; n = 4-5 cages/group. (F-J) The body weight (F) of male NTg mice was measured over the course of the experiment while fat (G) and lean mass (H) were determined at both the beginning and end of the experiment using EchoMRI and adiposity (I) was calculated; n = 6-10 mice/group. (J) Food consumption was measured in home cages at 10 months of age and normalized to body weight; n = 4-5 cages/group. (B-E, G-J) Statistics for the overall effects of diet, acarbose treatment (ACA), and the interaction represent the p-value from a 2-way ANOVA conducted separately for each time point (when applicable), with Sidak's post-test, $^{*}p < 0.05$, $^{**}p < 0.01$, $^{***}p < 0.001$, and $^{****}p < 0.0001$. Data represented as mean ± SEM.

### Figure S2

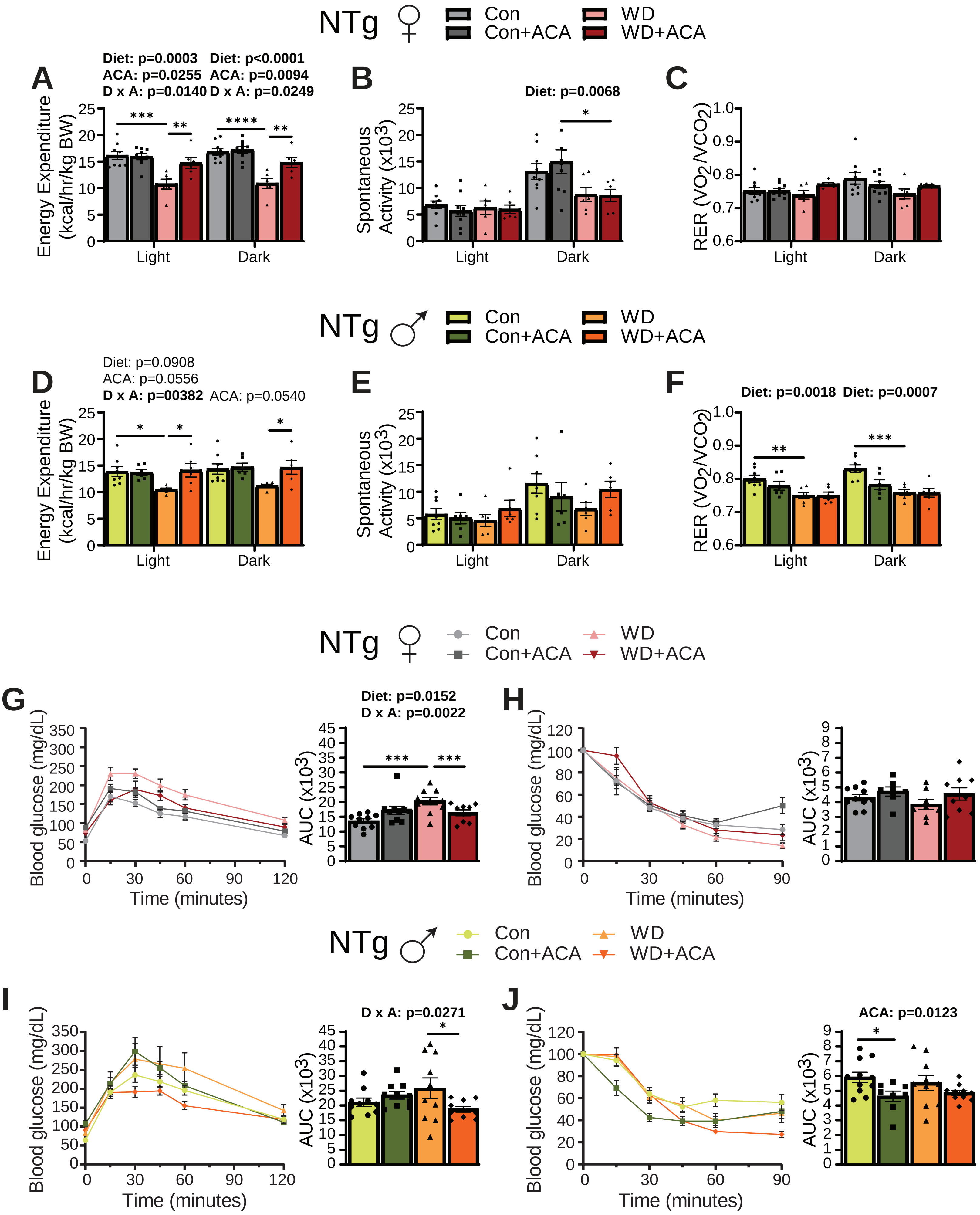
NTg ♀
Con
Con+ACA
WD
WD+ACA
A
Diet: p=0.0003
ACA: p=0.0255
D x A: p=0.0140
Diet: p<0.0001
ACA: p=0.0094
D x A: p=0.0249
Energy Expenditure (kcal/hr/kg BW)
Light
Dark
B
Diet: p=0.0068
Spontaneous Activity (x10³)
C
RER (VO2/VCO2)
NTg ♂
D
Diet: p=0.0908
ACA: p=0.0556
D x A: p=00382
ACA: p=0.0540
E
F
Diet: p=0.0018
Diet: p=0.0007
G
Diet: p=0.0152
D x A: p=0.0022
Blood glucose (mg/dL)
Time (minutes)
AUC (x10³)
H
I
D x A: p=0.0271
J
ACA: p=0.0123

**Supplementary Figure 2: The effects of WD on energy balance and glucose tolerance in NTg mice are opposed by acarbose.**

(A-F) Metabolic chambers were used to measure energy expenditure, spontaneous activity, and respiratory exchange ratio (RER) in female (A-C) and male (D-F) mice over a 24 hour period. (A, D) Energy expenditure normalized to body weight in females (A) and males (D). (B, E) Spontaneous activity of females (B) and males (E). (C, F) RER in females (C) and males (F). (G-H) Glucose (G) and insulin (H) tolerance tests were performed in female mice at 7 months of age, n = 9-10 mice/group. (I-J) Glucose (I) and insulin (J) tolerance tests were performed in male mice at 7 months of age. (A-F) n = 6-9, (G-J) n = 7-11 mice/group, statistics for the overall effects of diet, acarbose treatment (ACA), and the interaction represent the p-value from a 2-way ANOVA (conducted separately for light and dark periods in A-F), with Sidak's post-test, $*p < 0.05$, $**p < 0.01$, $***p < 0.001$, and $****p < 0.0001$. Data are represented as mean ± SEM.

### Figure S3

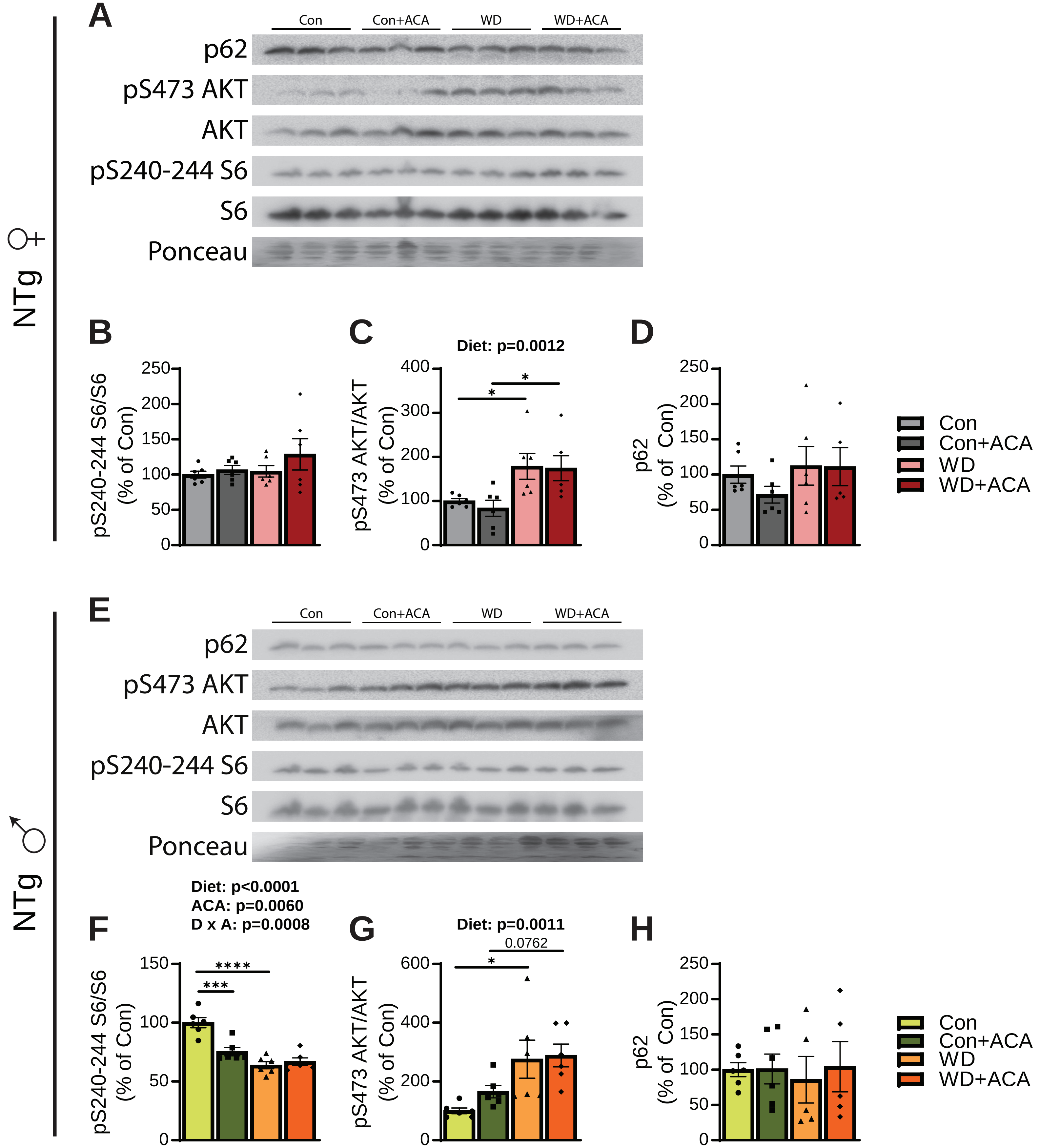

A
Con
Con+ACA
WD
WD+ACA
p62
pS473 AKT
AKT
pS240-244 S6
S6
Ponceau
NTg ♀
B
pS240-244 S6/S6 (% of Con)
C
Diet: p=0.0012
pS473 AKT/AKT (% of Con)
D
p62 (% of Con)
Con
Con+ACA
WD
WD+ACA
E
NTg ♂
F
Diet: p<0.0001
ACA: p=0.0060
D x A: p=0.0008
G
Diet: p=0.0011
0.0762
H

**Supplementary Figure 3: Sex-specific effect of WD and acarbose on mTOR signaling and autophagy in NTg mice.**

(A-H) mTOR signaling and autophagy were analyzed in the brains of 12-13 month old NTg mice via Western blotting of whole brain lysate. (A, E) Representative Western blots of female (A) and male (E) NTg mice. (B, F) Phosphorylation of S240/244 S6 relative to total S6 in female (B) and male (F) mice. (C, G) Phosphorylation of S473 AKT relative to total AKT in female (C) and male (G) mice. (D, H) Quantification of p62 relative to total protein (Ponceau) in female (D) and male (H) mice. (B-D, F-H) n = 5-6 mice/group; statistics for the overall effects of diet, acarbose treatment (ACA), and the interaction represent the p-value from a 2-way ANOVA, with Sidak's post-test, $*p < 0.05$ and $**p < 0.01$. Data represented as mean ± SEM

### Figure S4

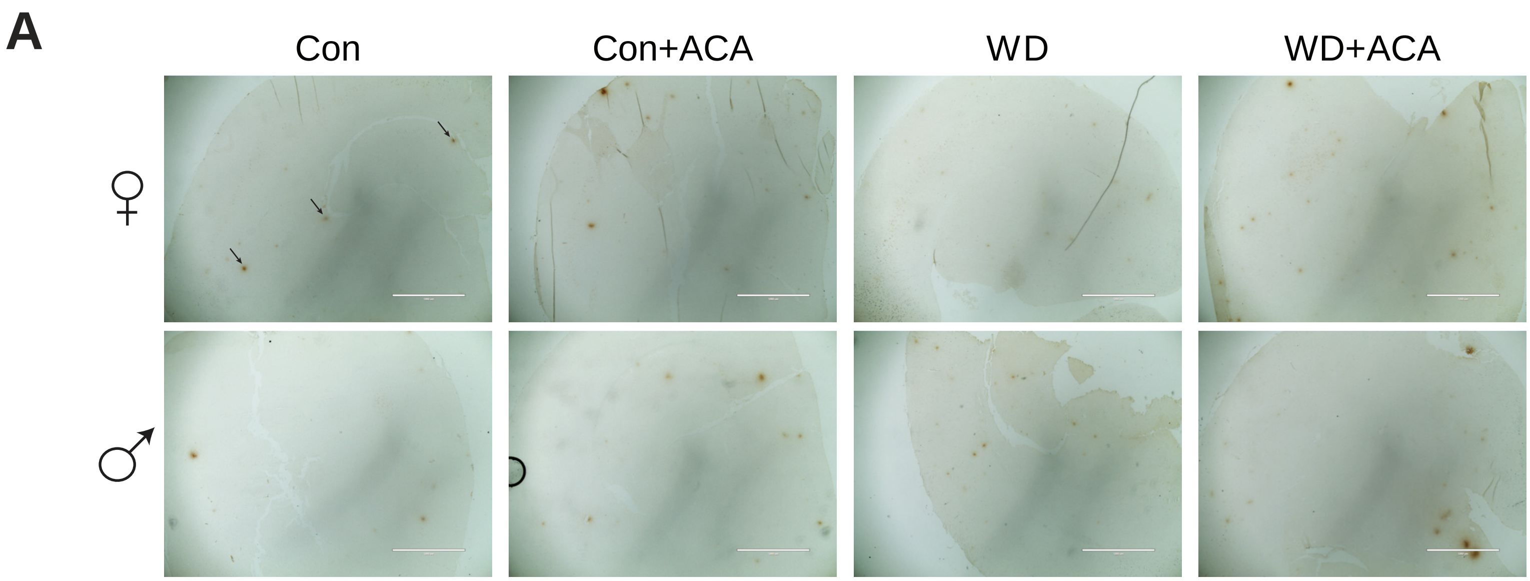

**Supplementary Figure 4: 3xTg mice have minimal aggregation of Aβ by 12 months of age.**

(A) Highest plaque load in the cortex and hippocampus of each sex/diet as determined by DAB staining specific to $A\beta_{1-16}$ (4x magnification).

### Figure S5

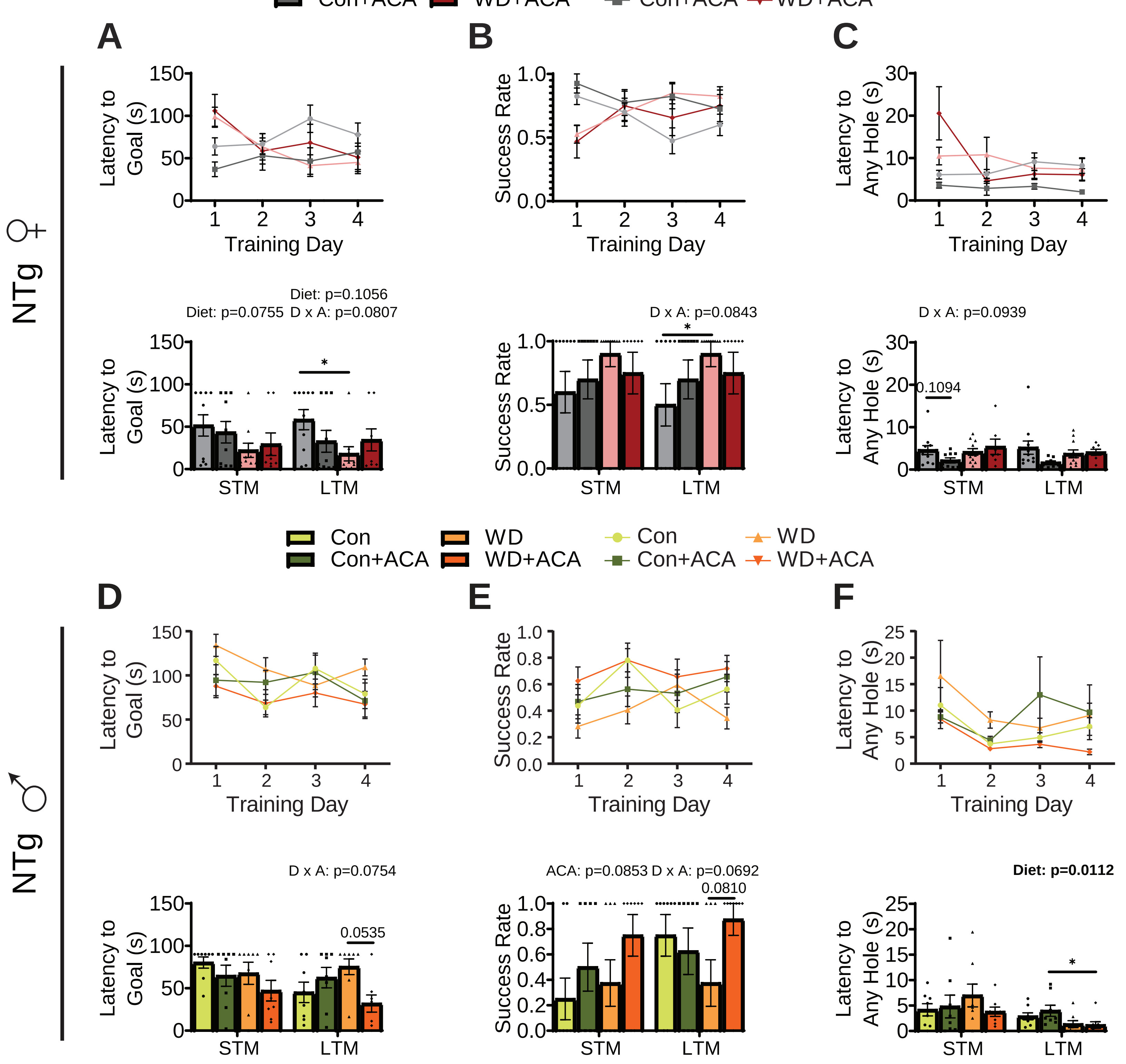

Con
Con+ACA
WD
WD+ACA
A
B
C
NTg ♀
Latency to Goal (s)
Success Rate
Latency to Any Hole (s)
Training Day
Diet: p=0.1056
Diet: p=0.0755 D x A: p=0.0807
D x A: p=0.0843
D x A: p=0.0939
0.1094
STM
LTM
D
E
F
NTg ♂
D x A: p=0.0754
0.0535
ACA: p=0.0853 D x A: p=0.0692
0.0810
Diet: p=0.0112

**Supplementary Figure 5: Spatial learning and memory in female NTg mice do not benefit from acarbose, while WD-fed NTg males show improvements in memory with acarbose.**

(A-F) Barnes maze was used to assess spatial learning and memory. (A, D) Latency to goal was measured in female (A) and male (D) mice. (B, E) Success rate for female (B) and male (E) mice. (C, F) Latency to any hole, a measure of anxiety, of female (C) and male (F) mice. (A-F) n= 8-10; statistics for the overall effects of diet, acarbose treatment (ACA), and the interaction represent the p-value from a 2-way ANOVA conducted for STM and LTM separately, with Sidak's post-test, *$p < 0.05$ and **$p < 0.01$. Data represented as mean ± SEM

### Figure S6

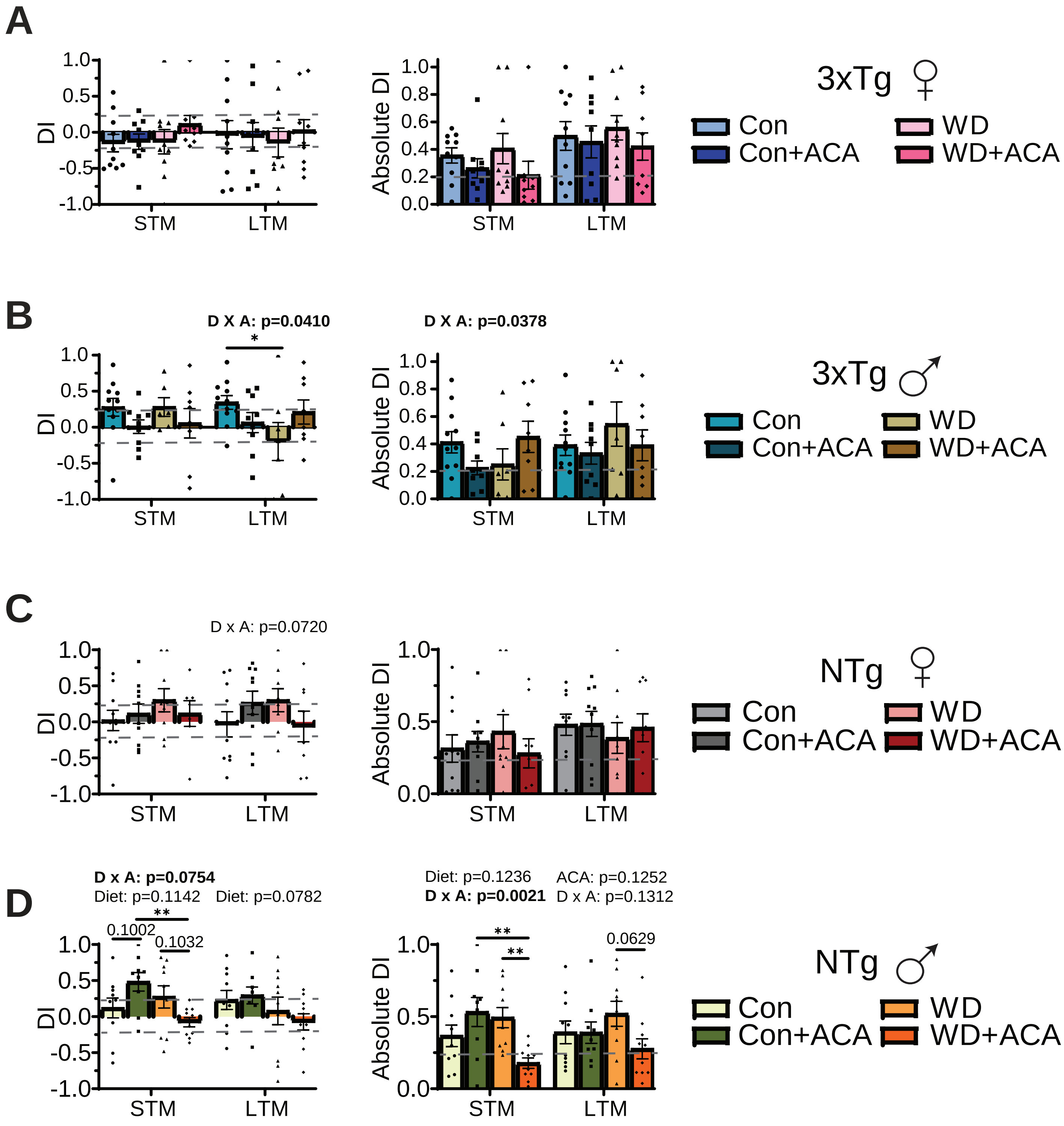
A
DI
Absolute DI
STM
LTM
3xTg ♀
Con
Con+ACA
WD
WD+ACA
B
D X A: p=0.0410
D X A: p=0.0378
*
3xTg ♂
C
D x A: p=0.0720
NTg ♀
D
D x A: p=0.0754
Diet: p=0.1142
Diet: p=0.0782
0.1002
**
0.1032
Diet: p=0.1236
D x A: p=0.0021
ACA: p=0.1252
D x A: p=0.1312
0.0629
NTg ♂

**Supplementary Figure 6: Effects of WD and acarbose on novel object recognition in 3xTg and NTg mice.**

(A-D) Novel object recognition (NOR) was utilized to assess recognition memory. Discrimination index (DI) and absolute DI were calculated for female 3xTg (A), male 3xTg (B), female NTg (C) and male NTg (D) mice. (A-D) n = 8-10; statistics for the overall effects of diet, acarbose treatment (ACA), and the interaction represent the p-value from a 2-way ANOVA conducted for STM and LTM separately, with Sidak's post-test, *$p < 0.05$ and **$p < 0.01$. Data represented as mean ± SEM

**Supplemental Tables**

**Supplementary Table 1:** Composition and calorie content for diets used in this study.

**Supplementary Table 2:** Antibodies used for Western blotting and immunohistochemistry.
